# Analysis of changes in inter-cellular communications during Alzheimer’s Disease pathogenesis reveals conserved changes in glutamatergic transmission in mice and humans

**DOI:** 10.1101/2024.04.30.591802

**Authors:** Katrina Bartas, May Hui, Wei Zhao, Desiree Macchia, Qing Nie, Kevin T. Beier

## Abstract

Analysis of system-wide cellular communication changes in Alzheimer’s disease (AD) has recently been enabled by single nucleus RNA sequencing (snRNA-seq) and new computational methods. Here, we combined these to analyze data from postmortem human tissue from the entorhinal cortex of AD patients and compared our findings to those from multiomic data from the 5xFAD amyloidogenic mouse model at two different time points. Using the cellular communication inference tool CellChat we found that disease-related changes were largely related to neuronal excitability as well as synaptic communication, with specific signaling pathways including BMP, EGF, and EPHA, and relatively poor conservation of glial-related changes during disease. Further analysis using the neuron-specific NeuronChat revealed changes relating to metabotropic glutamate receptors as well as neuronal adhesion molecules including neurexins and neuroligins. Our results that cellular processes relating to excitotoxicity are the best conserved between 5xFAD mice and AD suggest that excitotoxicity is the main common feature between pathogenesis in 5xFAD mice and AD patients.

## INTRODUCTION

Alzheimer’s disease (AD) is a complex disease characterized by systemic changes in brain function^1,2^. While the pathological hallmarks of AD are the accumulation of amyloid plaques and tau tangles in the brain, changes in cellular function occur in neurons and glia throughout the course of disease that contribute to the development of age-associated cognitive impairment^3,4^. Understanding the root molecular causes of disease is thus required for the development of effective therapeutics to combat the development of AD. Our ability to understand the molecular features of disease across the lifespan has been greatly accelerated by the recent development of single-cell and single-nucleus RNA sequencing (snRNA-seq) technologies. These methods have provided an atlas of gene expression throughout the course of disease, particularly in late-stage AD in brain regions such as the frontal cortex, entorhinal cortex (ENT), and the hippocampus^5–8^. However, a key limitation of these experiments is that they focus only on the cell-autonomous features of the disease; that is, they examine cells in isolation. While a reductionist approach is necessary to define the core features of disease, understanding how the molecular features of disease, as revealed by snRNA-seq, influence cellular communication is essential towards uncovering the root and progression of AD.

Fortunately, in recent years a variety of computational methods have been developed that enable the inference of cellular communication through coordinated changes in receptor-ligand interactions in different cell types. These include CellPhoneDB^9^ and CellChat^10^, among others. These methods have been widely used in a variety of tissue types to enrich single cell sequencing datasets and provide a wealth of information about modes of biological communication within tissues including in the AD brain. However, as these studies focused on glia and did not analyze data from neurons, they include only data from selected cell types^11–15^. While these methods’ utility in part is due to their ability to be generalized to different tissues, they also focus exclusively on receptor-ligand interactions that are encoded by proteins. While this is an acceptable definition for most tissue types, its use is more limited in the brain, where cells principally communicate through chemical transmission of small-molecule neurotransmitters, including glutamate and GABA, that are not encoded by proteins. To fill this gap, NeuronChat^16^ was developed to infer the intercellular communications mediated by small-molecule neurotransmitters. As such, if we want to understand how communication systems in the brain change during disease, an integrated understanding of how both general communication and neuron-specific modes of communication change during disease is required.

Here, we focus on the ENT, as it is a core region in the hippocampal circuit and thought to be a main contributor to AD-associated memory and cognitive deficits^17–20^. For these studies we used the 5xFAD mouse model, a commonly studied amyloidogenic model used in ∼10% of all AD studies employing rodent models^21,22^. We sequenced the ENT from 5xFAD and littermate control mice at 2 months of age, before memory deficits are apparent, and at 8 months of age, when behavioral deficits are widespread^22,23^. For comparison of gene expression changes in rodents to human postmortem brain tissue we utilized publicly available snRNA-seq data from the ENT of patients with AD and those without cognitive impairments. We also sequenced postmortem tissue from the ENT of a patient diagnosed with mild cognitive impairment (MCI) to provide a more comprehensive examination of AD progression in mice and human postmortem tissue across disease progression.

## METHODS

### Mouse and human tissue

Hemizygous 5xFAD mice (mutations in APP: Swedish, Florida, and London; mutations in PSEN1: M146L and L286V) on a C57BL/6J background (Jackson Laboratory, Stock #34848) were utilized. Wildtype littermates on the same background served as control animals. Female mice were used exclusively for all experiments. Mice were euthanized at either 2 or 8 months of age depending on the time point of the experiment. Mice were housed on a 12-hour light–dark cycle with food and water ad libitum. All procedures complied with the animal care standards set forth by the National Institute of Health and were approved by the University of California, Irvine’s Institutional Animal Care and Use Committee (IACUC). Human postmortem brain tissue samples were obtained from UC Irvine’s Alzheimer’s Disease Research Center (ADRC). The female donor was diagnosed with mild cognitive impairment (MCI), had tau tangle stage 2, plaque stage A, and was 85 years old (sample 28_18).

### Sample collection and single-nucleus sequencing in mice

Single-nucleus ATAC-seq and RNA-seq were conducted using the 10x Multiome kit. A merged sample comprising nuclei from two female 5xFAD mice and another merged sample from two female C57BL/6 wild-type mice, either at 2 months or 8 months of age, were used, for a total of eight mice.

Animals were anesthetized under isoflurane induction and underwent cardiac perfusion with cold 1xPBS to remove blood from the brain. Brain tissue was immediately extracted, and entorhinal cortex was freshly dissected out of each sample. From this point onwards, both mouse and human tissue were treated the same. To isolate the nuclei, tissue was homogenized using Lysis Buffer and manually disrupted using pestles, followed by a 4-minute incubation period. Samples were centrifuged at 500g for 5 minutes at 4°C, and the supernatant was discarded. Subsequently, 1 ml of Nuclei Wash and Resuspension buffer (NWR) was added and incubated for 5 minutes. The pellet was carefully resuspended using a P1000 pipette and centrifuged again at 500g for 5 minutes at 4°C. This wash-resuspension-centrifugation process was repeated twice more. After the third centrifugation, the supernatant was discarded, and 200μL of NWR buffer + 20μL 7AAD (1:100) was added and passed through a 40μm filter directly into a tube used for flow cytometry sorting. Cells were sorted at the UC Irvine Institute for Immunology Flow Cytometry Facility to remove debris and distinguish viable nuclei. Following sorting, the samples underwent a permeabilization process, which began with centrifugation at 500g for 5 minutes at 4°C. Subsequently, 100μL of 0.1X Lysis buffer was added, followed by resuspension and a 2-minute incubation. Next, 1mL of Washing Buffer was added, and the sample was centrifuged again for 5 minutes under the same conditions. Then, 500μL of Diluted Nuclei Buffer was added and carefully mixed with a pipette. Finally, the resulting sample was spun down and submitted to the UC Irvine’s Genomics Research and Technology Hub for further processing.

### Single-nucleus sequencing alignment and QC in mice

Alignment of the ATAC-seq and gene expression libraries to the mouse reference genome (mm10, v2020-A) was done using Cell Ranger ARC (v2.0.1, 10X Genomics). Alignment was performed on the High-Performance Community Computing Cluster (HPC3) at UC Irvine. Further processing was done in R largely using Signac^24^ (v1.7.0), which is integrated with Seurat^25^ (v4.3.0). Nuclei with the following QC metrics in the RNA-seq data were filtered out as they were likely doublets, debris, or damaged: over 20 percent of reads aligned to mitochondrial genes, more than 70,000 or fewer than 200 UMIs, or more than 10,000 or fewer than 75 unique genes detected (Figure S1A-B). Nuclei with the following metrics were further filtered out from the ATAC-seq data, to prioritize only high-quality nuclei: fewer than 10 percent reads in peaks, nucleosome signal score of greater than 3, transcription start site enrichment of less than 2, over 5 percent of reads being in a blacklist region, or counts in peaks of under 1,000 or over 70,000. Following QC, there were a total of 35,188 high quality nuclei remaining: 8,286 from 2-month-old C57BL/6 mice, 15,126 from 2-month-old 5xFAD mice, 6,431 from 8-month-old C57BL/6 mice, and 5,345 from 8-month-old 5xFAD mice.

### Single-nucleus sequencing alignment and QC, human postmortem samples

Previously published snRNA-seq data collected from the ENT of AD and non-cognitively impaired, or control, patients was downloaded from the gene expression omnibus^5,26^ and re-processed using Seurat. Only female patient data was included for consistency with our other dataset from exclusively female mice. From these two previously published datasets, data from 8 patients were used: 4 female patients with AD, and 4 female patients without cognitive impairments. The additional female MCI patient sample was sequenced at UCI’s GRT Hub and resulting fastqs aligned to the NCBI Homo Sapiens Annotation Release 109 using the Parse Biosciences computational pipeline (v1.0.5p). The resulting gene by cell counts matrices were QCed individually by sample and subsequently integrated using Seurat’s IntegrateData function with default parameters.

### Single-nucleus sequencing integration, dimensionality reduction, and clustering, mice

To integrate 5xFAD and control mouse brain transcriptomic data, sctransform-based normalization was used to normalize, scale, and find variable features within each dataset^27^. Mitochondrial mapping percentage, a potential confounder, was regressed out, and the Gamma-Poisson Generalized Linear Model method^28^ was used, otherwise default sctransform parameters were used. 3,000 features that varied the most across all datasets were selected as integration features, and the datasets were integrated using the IntegrateData command in Seurat, with anchor features being set to our list of 3,000 features, the normalize method specified as sctransform, and the other parameters set to their default values. This resulted in one Signac object containing all mouse data, including the 2-month-old and 8-month-old 5xFAD and WT mice. After integration, the standard Seurat workflow for dimensionality reduction, visualization, and clustering was run sequentially using the following commands: RunPCA, RunUMAP, FindNeighbors, and FindClusters. For RunUMAP and FindNeighbors the first 30 principal components were used, and for RunUMAP the number of nearest neighbors parameter was set to 30. The resolution parameter for FindClusters was set to 1.0.

### Annotation of cell types

Cell types in our mouse data were predicted by transferring annotation labels from the Allen Brain Atlas’ transcriptomic data^29,30^ to our datasets, and this predicted annotation was manually checked and refined using previously established marker genes^31–33^. Oligodendrocyte markers used were *Mobp*, *Il33* and *Mog*, OPC markers were *Cspg4*, *Tnr*, and *Pdgfra*, astrocyte markers were *Aqp4*, *Slc1a2*, and *Gja1*, microglia markers were *Csf1r*, *Ctss*, and *C1qb*, GABAergic neuron markers were *Gad1* and *Gad2*, glutamatergic neuron markers were *Slc17a7* and *Slc17a6*, and endothelial cell markers were *Bsg*, *Flt1*, and *Vwf*. The same marker genes were used to manually annotate the data from postmortem human samples.

### snATAC-seq data processing, mice

Following integration and annotation, we re-processed the ATAC data. As peaks had previously been called on each sample individually by Cell Ranger ARC, they needed to be redefined so that common peaks would be identified across all mouse brain samples. We also aimed to call peaks by cell type to identify peaks in rare cell types that were potentially missed in aggregate. On the integrated data we ran the CallPeaks function in Signac, which uses MACS2 to call peaks^34^. We performed this peak calling by cell type, annotated according to the RNA-seq data, and these peak sets were then combined. Peaks found to be in genomic blacklist regions or on nonstandard chromosomes were removed.

To visualize the ATAC data we performed the standard Signac workflow with the following commands: RunTFIDF, FindTopFeatures, RunSVD, and RunUMAP. We then combined this representation of the data with the transcriptomic data visualization. This representation of both RNA and ATAC data was created using the FindMultiModalNeighbors function in Signac, followed by running UMAP on the weighted nearest neighbor (WNN) graph.

### Differential gene expression analysis

Before analysis, counts were normalized using the NormalizeData command in Seurat. Differential gene expression was performed on each cell type cluster using the R package presto^35^. Genes were considered significantly different if the adjusted p-value (Bonferroni correction) was less than 0.05. Comparisons were made between 5xFAD and WT mice at 2 months of age, 5xFAD and WT mice at 8 months of age, AD and cognitively non-impaired patient samples, and MCI and cognitively non-impaired patient samples.

### CellChat and NeuronChat analysis

CellChat (v1.6.1)^10^ and NeuronChat (v1.0.0)^16^ were used to infer communication between cell types. First, CellChat and NeuronChat’s standard workflows were run on each condition within the snRNA-seq data individually: 2-month-old 5xFAD mice, 2-month-old WT mice, 8-month-old 5xFAD mice, and 8-month-old WT mice. Separately, human samples were also put through this pipeline: AD, MCI, and control. Comparative analyses were then performed between 5xFAD and WT mice at 2 months of age and 8 months of age. AD and cognitively normal patient samples, and MCI and cognitively normal patient samples were also compared.

### hdWGCNA, mouse and human comparison

To detect modules of genes that co-vary in a cell type-specific manner, high dimensional weighted gene co-expression network analysis (hdWGCNA) was performed using the hdWGCNA package^36^. hdWGCNA adapts the well-established WGCNA method^37^ for snRNA-seq data. This analysis was performed on cell types where there were at least 1,000 nuclei in the mouse dataset: microglia, oligodendrocytes, OPCs, astrocytes, GABAergic neurons, and glutamatergic neurons. Modules were detected in each cluster within the mouse snRNA-seq data, projected onto the human data, and then module preservation and module quality were evaluated. Modules found to be of high quality and highly preserved across species (zsummary.qual >10 and zsummary.pres > 10) were further investigated. Differential module eigengene analysis was performed to analyze differences in expression of each module between control and disease groups.

This analysis was performed, and results visualized using EnrichR (v3.178)^38,39^ and hdWGCNA functions RunEnrichr, GetEnrichrTable, and EnrichrBarPlot sequentially. Enrichment analysis was done using the 2021 versions of the Gene Ontology databases^40,41^.

### DIRECT-NET analysis and visualization, mice

DIRECT-NET^42^ was applied to the mouse 10X Genomics Multiome data to detect regulatory links specific to either 5xFAD or WT mice at 2 and 8 months of age and to construct gene regulatory networks implicated in AD. Promoters for each gene in the mouse genome were defined to be the 1,000 basepairs before the transcription start site of a gene. The DIRECT-NET tutorial was then followed, with analyses being applied to each genotype and time point individually first and then compared. First, cis-regulatory elements (CREs) were inferred following the DIRECT-NET tutorial for genes of interest based on previous analyses, including *Gria4*, *Grin2b*, and *Grm1*. Links were detected by genotype by subsetting the data and then running the Run_DIRECT_NET function on each genotype at each age. Signac’s coverage plot function was used to visualize these links. Only high-confidence (HC) links were plotted, with links specific to 5xFAD mice highlighted in pink, links specific to control mice highlighted in blue, and links apparent in both genotypes highlighted in purple.

We then proceeded to construct gene regulatory networks based on genes upregulated in 5xFAD mice for both the 2- and 8-month time points. Following the DIRECT-NET tutorial, we used presto^43^ to determine genes that were upregulated in 5xFAD mice according to auROC analyses (auc > 0.5) of snRNA-seq data. We excluded genes corresponding to ribosomal proteins and those without known functions, then evaluated which of the remaining genes overlapped with regions of the genome that were more accessible in 5xFAD mice based on the accompanying snATAC-seq data. CREs relating to both upregulated and accessible genes were then used to generate links to known transcription factors using the generate_peak_TF_links command in DIRECT-NET. These links were then constructed into a network and visualized using the Python package networkx^44^.

## RESULTS

### Sample selection and quality control for mouse and human AD samples

One significant advantage of using rodent models of AD is that AD pathology can be studied across the lifespan, including at pre-symptomatic time points. We chose two time points for this study: 2 months, a pre-symptomatic time point when plaques begin to form in select regions in the brain, and 8 months, after significant plaque accumulation has occurred and cognitive deficits have emerged^22,23^. We used female mice only, as female 5xFAD mice show more rapid pathogenesis than males^45–47^. We similarly only evaluated tissue samples from female postmortem brain tissue, as women have an increased risk of developing AD compared to men^48^. We performed a multiomic assessment (single nucleus RNA-seq plus single nucleus ATAC-seq) of cells from the ENT of 5xFAD and age-matched controls from each age group (Figure 1A). After alignment and quality control assessments of data, we had 23,412 cells for the 2-month time point and 11,776 cells for the 8-month time point. All expected cell types were captured in each dataset, including excitatory neurons, inhibitory neurons, astrocytes, microglia, oligodendrocytes, and oligodendrocyte precursor cells (OPCs; Figure 1B-C). Cell types were similarly represented across time points and genotypes (Figure S1C-E). To confirm that our integrated multiomic strategy worked as expected, we assessed the correspondence between the snRNA-seq and snATAC-seq datasets. Marker genes such as *Slc17a7*, *Mog*, and *C1qb* were more accessible and more highly expressed in glutamatergic neurons, oligodendrocytes, and microglia, respectively, than in other cell types (Figure 1C-F), consistent with known gene expression patterns. Accessibility of other marker genes for each cell type are shown in coverage plots in Figure S3B-Q.

**Figure 1:**
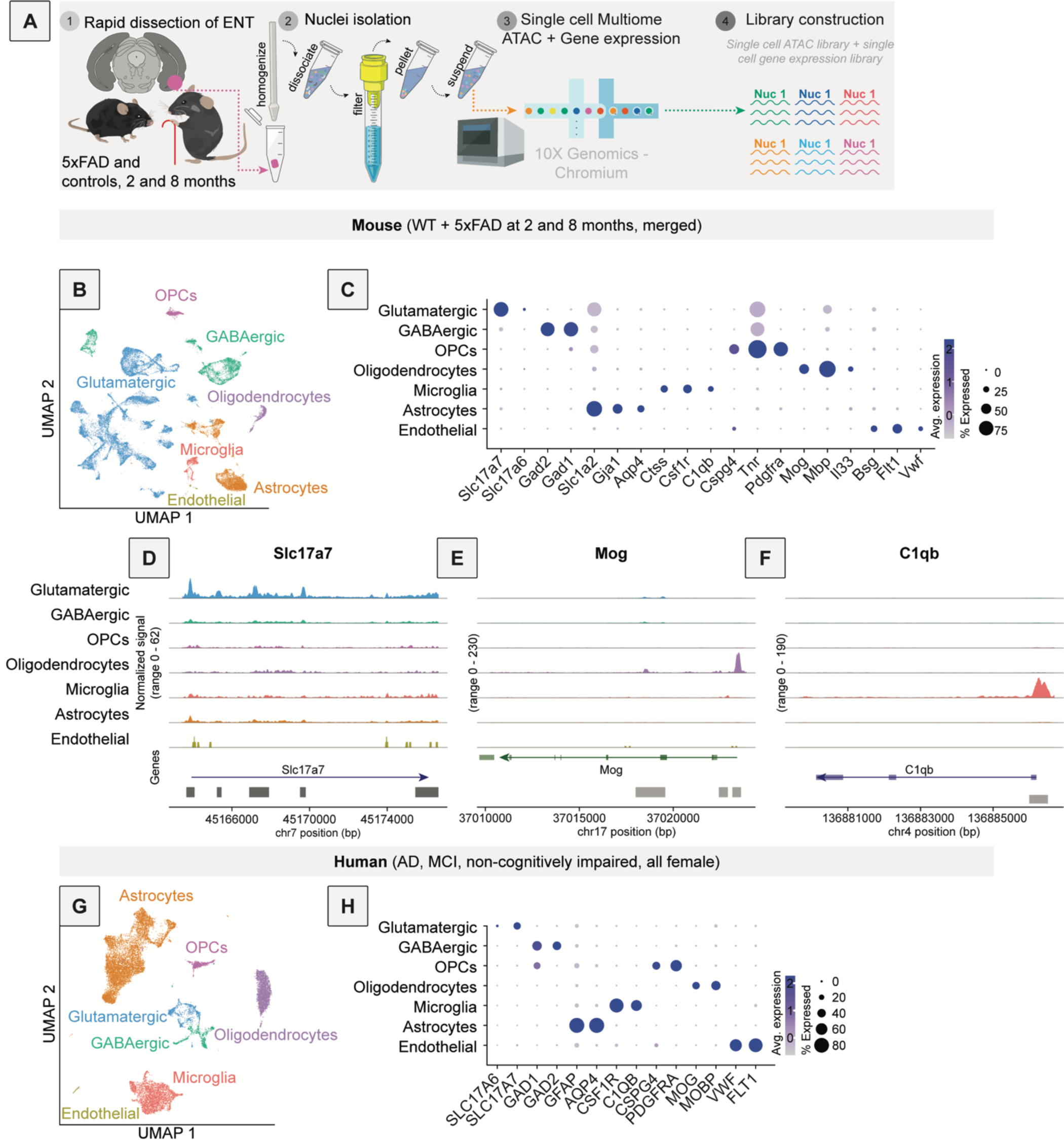
Characterization of single cell sequencing data from mouse and human postmortem brain samples. **A**, Diagram of experimental workflow for snRNA-seq and snATAC-seq. The ENT of mice aged 2 months and 8 months, both 5xFAD and control, was dissected, nuclei were isolated, and sequencing performed using the Single Cell Multiome ATAC + Gene Expression kit from 10x Genomics. **B**, Single-nucleus data for mice aged 2 months and 8 months, both 5xFAD and control, represented in 2D UMAP (WNN) with general cell type annotations. **C**, Dotplot showing expression of known marker genes in each cell type in the mouse snRNA-seq portion of the multiome data. **D-F**, Coverage plots showing known marker gene accessibility as captured by the snATAC-seq portion of the multiome data. Chromatin accessibility was reflective of gene expression (i.e., the area near the transcription start site is more open in genes that are more highly expressed in defined cell types). Shown are coverage plots for *Slc17a7* (**D**), a marker for glutamatergic neurons; *Mog* (**E**), a marker for oligodendrocytes; and *C1qb* (**F**), a marker for microglia. **G**, UMAP of merged, re-processed, and annotated snRNA-seq data from postmortem tissue samples obtained from AD, MCI, and non-cognitively impaired human brains. This is a combination of previously published datasets from female donors^5,26^, and one newly sequenced tissue sample. **H**, Dotplot showing expression of known marker genes in the annotated human snRNA-seq data.

To explore the diversity of neuron types identified, we assessed the number of distinct subtypes of glutamatergic and GABAergic neurons within our mouse dataset. We performed unsupervised clustering using the smart local moving (SLM) algorithm and found 5 inhibitory (In) and 12 excitatory (Ex) neuronal subclusters with distinct transcriptional profiles (Figure S2A). We also transferred labels from a well-annotated dataset from the Allen Brain Institute that enabled definition of each cell type’s layer in the brain and assessed gene expression differences (Figure S2B-C). Using this annotation method, we identified the following cell types: GABAergic subcluster 1 (In 1) was defined by expression of the vasal intestinal peptide (*Vip*), In 2 by synuclein gamma (*Sncg*), In 3 by lysosomal-associated membrane protein 5 (*Lamp5*), In 4 by parvalbumin (*Pvalb*), and In 5 by somatostatin (*Sst*). Excitatory neurons were classified by layer (depth in cortex, L2-6) and connectivity patterns (intratelencephalic, IT; corticothalamic, CT; pyramidal tract, PT; near projecting, NP).

To assess how our mouse results compared with those from human postmortem brain tissue, we utilized data from previously published studies from ENT tissue in humans with and without AD. We also performed snRNA-seq of postmortem brain tissue from one female patient with mild cognitive impairment (MCI) to acquire data from an earlier stage of disease. After merging and processing the human data, we observed the same cell types as we found in our mouse data, though in different quantitative proportions (Figure 1G). Annotation of cell types in the human data was based on the same marker genes as used in the mouse data (Figure 1H, 1C). As we identified fewer neurons in the human samples, we did not perform further annotation of neurons beyond inhibitory and excitatory.

### Differential gene expression and high dimensional weighted gene correlation network analysis (hdWGCNA) indicate a greater conservation of neuron-related gene expression changes relative to glial-related gene expression changes between mouse models and human AD

We next explored differences in gene expression between 5xFAD versus control mice, and human AD/MCI postmortem samples versus samples from non-cognitively impaired human postmortem brain tissue. Results of differential gene expression analysis for each cell type and each of the four comparisons (5xFAD versus control at 2 months, 5xFAD versus control at 8 months, MCI versus control, and AD versus control) including fold-change and adjusted p-value are reported in Supplementary Table 1.

As a benchmark, we first focused on several genes known to be associated with disease-associated astrocytes (DAAs) that emerge during the development of AD: Clusterin (*Clu*), Apolipoprotein E (*Apoe*), and Cystatin C (*Cst3*) ^49,50^. DAAs have previously been identified in the hippocampus and cortex of both human AD patients and 5xFAD mice^49,50^. Genes upregulated in DAAs are associated with amyloid clearance, aging, and the complement system^51^. Unexpectedly, at 2 months of age in 5xFAD mice, DAA genes were downregulated compared to controls (Figure 2A). Interestingly, we see a very similar downregulation of *APOE* and *CST3* in the human MCI brain (Figure 2B). However, in 5xFAD mice at 8 months of age the DAA genes *Apoe*, *Clu,* and *Cst3* were significantly upregulated, consistent with previous reports in mice and humans that these genes are enriched in DAAs or upregulated in AD in subclusters of astrocytes (Figure 2C)^49,52^. We observed a significant upregulation of each of these genes in the human AD brain (Figure 2D). These data together indicate that gene expression patterns occurring in the earliest stages of disease may not mirror those that occur at later stages, and in some cases may trend in the opposite direction. This in turn may indicate that gene expression changes during AD pathogenesis may not progress in a linear fashion and could be dynamically regulated during disease.

**Figure 2:**
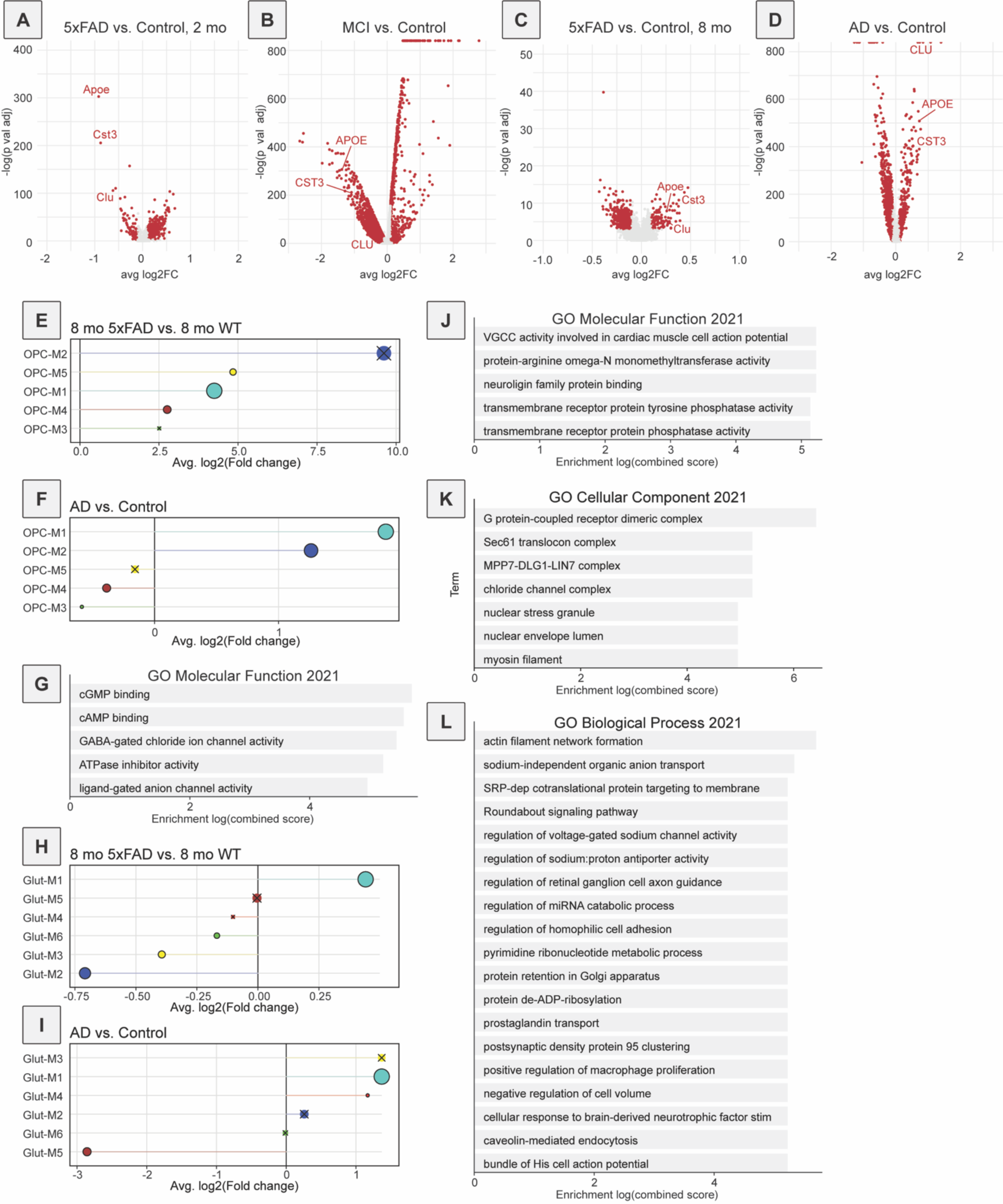
Differential gene expression and conserved modules of gene expression changes from mouse and human data using hdWGCNA. **A**, Differential gene expression in astrocytes for 5xFAD versus control mice at 2 months of age. A positive log-fold change indicates that a gene is upregulated in the 5xFAD genotype. **B**, Differential gene expression in astrocytes for human MCI vs non-cognitively impaired controls. A positive log-fold change indicates that a gene is upregulated in MCI. **C**, Differential gene expression in astrocytes for 5xFAD versus control mice at 8 months of age. A positive log-fold change indicates that a gene is upregulated in the 5xFAD genotype. **D**, Differential gene expression in astrocytes for human AD vs non-cognitively impaired controls. A positive log-fold change indicates that a gene is upregulated in AD. **E**, DME analysis of OPC modules in 8-month-old 5xFAD mice. OPC M1 was significantly upregulated in 5xFAD mice at 8 months. If a module was not significantly different between conditions, an ‘x’ indicates an adjusted p-value of > 0.05. **F**, DME analysis of OPC modules in human AD compared to non-cognitively impaired individuals. The gene expression module OPC M1 was significantly upregulated in AD. **G**, GO analysis of genes within OPC M1, from molecular function database. cGMP and cAMP binding were among the most prominent terms, as were those related to ion channels. **H**, DME analysis of glutamatergic neuron modules in 8-month-old 5xFAD mice. Glut M1 was upregulated in 5xFAD mice at 8 months. **I**, DME analysis of glutamatergic neuron modules in human AD. Glut M1 was upregulated in human AD. **J**, GO analysis of genes within Glut M1, molecular function database. Two of the top terms relate to voltage-gated calcium channels, which control cellular excitability, and neuroligin-mediated cellular interactions. **K**, GO analysis of genes within Glut M1, cellular component database. G-protein coupled receptors, which can mediate a variety of cellular processes including excitability, and chloride channels were among the top terms. **L**, GO analysis of genes within Glut M1, biological process database. Highlights include genes relating to intrinsic neuronal excitability and cell adhesion.

While assessing changes in the expression of individual genes is useful to assess specific changes, a more holistic approach may better represent the overall patterns of gene expression changes during disease progression. Previously, weighted gene co-expression network analysis (WGCNA^37^) was applied to bulk RNA-seq data to detect such gene networks, which allowed for further analysis and comparison of these gene modules. This approach was recently modified for use with single-cell data (hdWGCNA^36^). Here we applied hdWGCNA to each cell population in the mouse dataset (oligodendrocytes, microglia, OPCs, astrocytes, GABAergic neurons, and glutamatergic neurons). We then projected these cell type-specific mouse gene modules onto the corresponding human gene expression data and quantified module preservation across species and module quality^53^. Only high-quality modules, meaning reproducible and robust, and highly preserved modules with a similar network architecture in both species were further examined. This quality and preservation analysis was done to focus our efforts on modules whose expression change was not due to chance, and was not unique to the mouse model to increase the likelihood that our results are relevant to human AD.

Interestingly, we found that in microglia and astrocytes, two cell types commonly studied in the context of AD in rodent models including 5xFAD^54–57^, no modules were both highly preserved and of high quality (Figure S4A-B), indicating that patterns of gene expression changes were distinct in 5xFAD mice and human AD for these cell types. In oligodendrocytes just one module, Oligo-M3, was both highly preserved and of high quality (Figure S4C). To evaluate whether expression of genes in this module was significantly different between genotypes, we used differential module eigengene (DME) analysis implemented in hdWGCNA and found that the module expression was significantly upregulated in 2-month-old 5xFAD mice and in postmortem brain samples from both human MCI and AD compared to samples from non-cognitively impaired patients (Figure S4D-G). In 8-month-old 5xFAD mice a similar effect was seen but it was not statistically significant (Figure S4E). Similarly, in GABAergic neurons one module, GABA-M3, was both highly preserved and of high quality (Figure S5A). This module was not significantly differentially expressed between samples from 5xFAD and control mice at either age, nor postmortem brain samples from AD or MCI patients and controls (Figure S5B-E), suggesting that the differences in expression were relatively mild. In OPCs, one module, OPC-M1, was both highly preserved and of high quality (Figure S5F). The downregulation of genes in this module was not significant at 2 months (Figure S5G), however, this module was downregulated in MCI (Figure S5H) but upregulated in human AD and in 5xFAD mice (Figure 2E-F). We further investigated what genes were expressed in OPC-M1 and to what pathways they may contribute. OPCs primarily differentiate into oligodendrocytes, but there is also evidence that they can become neurons or astrocytes under certain conditions^58–61^. OPCs are involved in biological processes including myelination^62^ as well as synaptic remodeling and pruning^63–65^. Based on gene ontology (GO) analysis, genes in OPC-M1 were most highly associated with cGMP and cAMP binding (Figure 2G). cGMP and cAMP are both well-known small molecule second messengers for a variety of signal transduction pathways and have both been shown to modulate synaptic plasticity^66–68^. Additionally, it has previously been suggested that cAMP and cGMP alter metabolism of amyloid-β or stimulate its production^69–71^, which is especially relevant given amyloid-β’s abundance in the 5xFAD mouse brain.

Overall, the best module quality and conservation was observed in glutamatergic neurons, where we observed 2 modules, GluT-M1 and GluT-M3, that were both highly preserved and of high quality (Figure S5I). DME analysis indicated that only GluT-M1 was significantly differentially expressed – upregulated – in both 5xFAD mice at both time points and postmortem human AD brains relative to their respective controls (Figure 2H-I, Figure S5J-K). Differences were not significant in MCI. GO analysis of this gene module highlighted biological processes including voltage-gated calcium channel activity (Figure 2J), G protein-coupled receptors (GPCRs; Figure 2K), and actin filament network formation (Figure 2L), all processes that are important in synaptic regulation^72–74^. Interestingly, though the GABA-M3 module noted previously was not significantly differentially expressed, GO analysis of GABA-M3 also yielded terms associated with calcium transport and glutamate receptors (Figure S5L), indicating a common set of neuronal changes relating to cellular excitability. Together, these results suggest that gene expression patterns related to neuron-specific signaling processes, including synaptic modulation by OPCs, intrinsic cellular excitability, and glutamatergic signaling are better preserved between 5xFAD mice and human AD brains than modules specific to glial cells, and that these neuronal processes may play a central role in disease pathophysiology.

### Changes in inter-cellular communication networks during AD development

While single cell sequencing data enable the study of gene expression and gene module regulation within defined cell types, one key limitation is that these results only focus on cell-intrinsic changes. This provides only a partial picture of disease, as the brain consists of numerous cell types that operate in a dynamic fashion during both health and disease, and appropriate inter-cellular communication is necessary for proper brain function. For example, as glial cells play critical roles in synaptic function and remodeling which occur in neurons, focusing only on the transcriptome of glial cells in isolation, and not considering how glial changes impact communication with neurons, may mask their true importance in disease. To gain a deeper understanding of the evolution of inter-cellular interactions within the ENT, we utilized the computational package CellChat^75^. This method infers changes in cell-cell communication networks based on known cellular communication pathways. CellChat determines potentially important signaling pathways by identifying in which cell populations known ligands and corresponding receptor genes are expressed, including multimeric proteins, as well as other proteins that may be involved. Based on expression of these genes, we can infer modes of inter-cellular communication, which can be visualized as a directed network where nodes are cell types and edges are connectivity probabilities. These networks can inform us of likely communication pathways between identified cell types, and how the strength of these predicted interactions change across conditions and/or disease.

We first assessed the overall inferred level of incoming and outgoing signaling for each cell type, based on snRNA-seq data. In both 2- and 8-month-old control mice, the majority of inter-cellular signaling – both outgoing and incoming – was predicted to either arise from or be received by OPCs or neurons, with microglia and endothelial cells predicted to contribute the least to inter-cellular signaling (Figure 3A-B). Notably, only relatively small differences were detected in 5xFAD mice relative to controls when comparing data from 2- and 8-month-old animals, primarily in astrocytes and oligodendrocytes (Figure 3A-B), indicating only subtle changes in inferred inter-cellular communication occur over time in 5xFAD mice. In contrast, while we observed similar elevations in predicted signaling in astrocytes and oligodendrocytes in human MCI and AD tissue samples, changes in predicted signaling in glutamate and GABA cells was much more pronounced, with more outgoing and incoming signaling predicted in each cell type in MCI and AD than controls (Figure 3C-D, S6A). These data together indicate that 5xFAD mice may exhibit similar patterns of signaling changes as in human AD, though to a lesser extent (Figure 3A-C, S6A-B).

**Figure 3:**
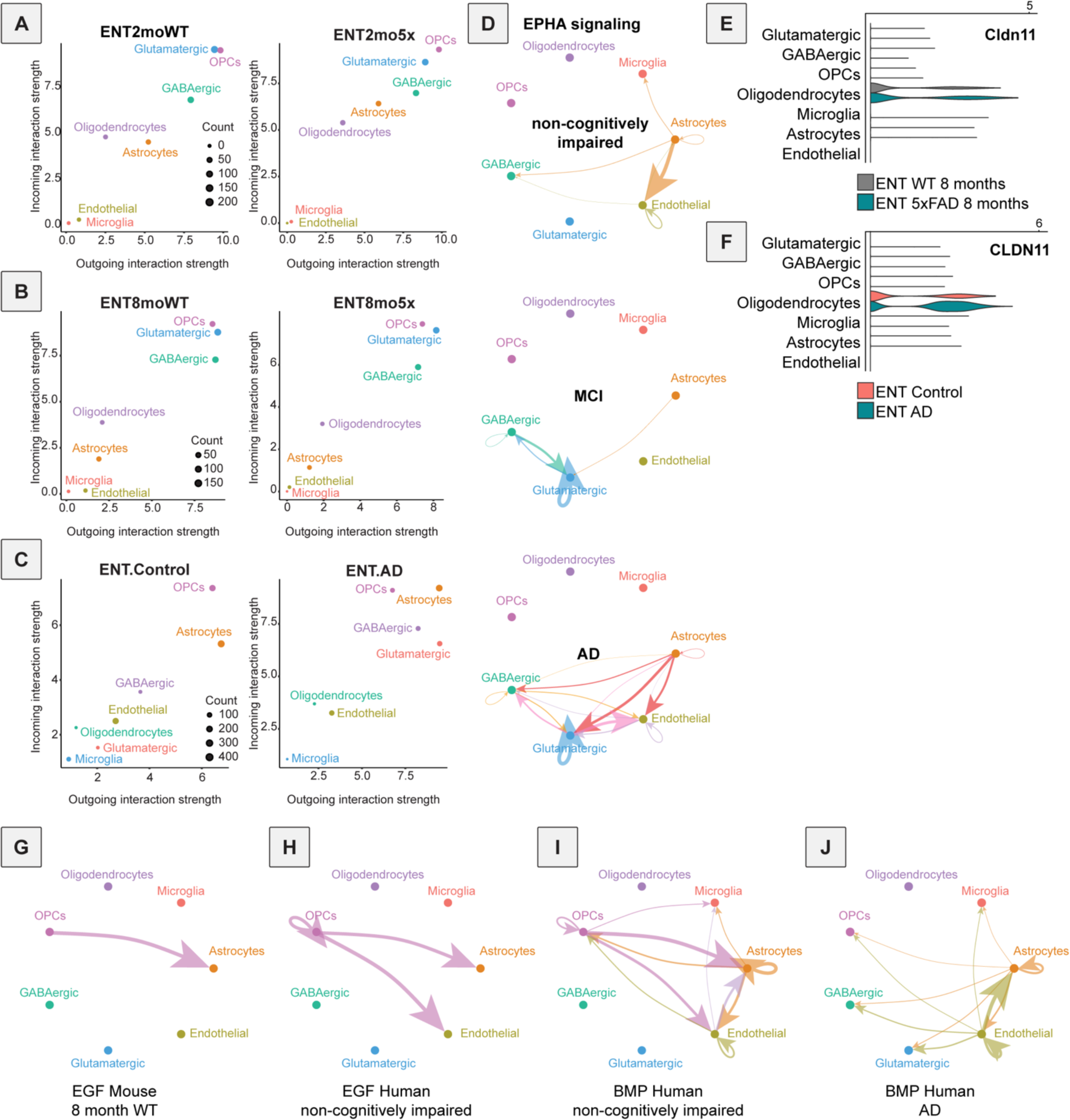
Application of CellChat reveals common and species-specific changes in cellular communication in mouse and postmortem human brain tissue. **A**, Predicted outgoing and incoming cellular signaling with CellChat for each cell type in the mouse ENT at 2 months of age for control mice (left) and 5xFAD mice (right). **B**, Predicted outgoing and incoming cellular signaling with CellChat for each cell type in the mouse ENT at 8 months of age for control mice (left) and 5xFAD mice (right). **C**, Predicted outgoing and incoming cellular signaling with CellChat for each cell type in the human ENT for patients without cognitive impairment (left) and patients with AD (right). **D**, CellChat signaling inferred for EPHA in patients (from top to bottom) without cognitive impairment, with MCI, and with AD. **E**, Violin plot of *Cldn11* expression at 8 months of age in mice. **F**, Violin plot of *Cldn11* expression in AD tissue samples, and from individuals without cognitive impairment. **G**, CellChat signaling inferred for EGF in mouse, control, 8 months of age. **H**, CellChat signaling inferred for EGF from brain tissue from donors without cognitive impairment. **I**, CellChat signaling inferred for BMP in samples from patients without cognitive impairment. **J**, CellChat signaling inferred for BMP from brain tissue from donors with AD.

We next identified molecular signaling pathways that showed similar or different changes between 5xFAD mice and human AD samples (Figure S6B). Pathways that exhibited the most similar disease-related changes across species were Ephrin-A (EPHA), Claudin (CLDN), Epidermal Growth Factor (EGF), and Bone morphogenetic protein (BMP). Predicted signaling through EPHA pathways was highly conserved between mice and humans across time points, and there was a higher level of predicted signaling in the EPHA pathway in 5xFAD mice at both timepoints as well as human MCI and AD compared to controls (Figure 3D, S6B-D). Glutamatergic neurons showed the largest increase in predicted EPHA signaling in the disease state relative to controls, an effect that was more pronounced in human MCI and AD than 5xFAD mice (Figure 3D, S6B-D). While the EPHA signaling pathway includes the genes *Epha1*-*8* and *Efna1*-*5*, this elevation in inferred signaling was mostly driven by expression changes in *Epha4* and *Epha7*, which were significantly upregulated in glutamatergic neurons in AD compared to controls (*EPHA4*, logFC = 0.695, adjusted p-value = 0.000557; *EPHA7*, logFC = 0.386, adjusted p-value = 0.00468, Supplementary Table 1). Interestingly, a loss of EphA4 was previously shown to improve aspects of social memory^76^ and synaptic plasticity in the hippocampus^77^, suggesting that an upregulation in EphA4, as observed here in AD-related disease, may impair properties of neuronal signaling. Interestingly, other studies have also linked EphA4 to amyloid-β, though the nature of this link is not clear and may be bidirectional. For example, while some studies suggested that Epha4 regulates amyloid-β^78^, others indicated that amyloid-β regulates *Epha4*^77^. While our results do not provide an answer to this discrepancy, they support the observation that as amyloid-β accumulates, *Epha4* expression increases.

Notably, while glutamatergic neurons showed the largest predicted differences in EPHA signaling, the *Epha3*, *Epha4*, *Epha5*, and *Epha7* genes were also present in the high-quality and highly preserved OPC-M1 discussed previously (Supplementary Table 2). In addition, the two GO terms previously mentioned as highly associated with OPC-M1 – cAMP and cGMP signaling – are engaged in the EPHA signaling pathway^79^, and are upregulated, consistent with an elevation in signaling through EPHA. Therefore, an upregulation in EPHA signaling may impact multiple cell types and contribute to neuronal dysfunction during the development of AD-related disease.

In the CLDN signaling pathway, only one gene was implicated, *Cldn11*, which was significantly upregulated in both 8-month-old 5xFAD mice and postmortem human AD tissue samples (Figure 3E-F). We found that the upregulation of *Cldn11* in oligodendrocytes is opposite of that found in a previous study performed in human postmortem AD samples, where a downregulation was observed^80^. Notably, that study was performed using tissue from the precuneus region in the brain rather than the ENT, and therefore, one possibility is that changes may be brain region-dependent. Supporting this interpretation, another study using the amyloidogenic APP-NL-G-F^81^ mice found that a gene module containing *Cldn11* was either upregulated or downregulated depending on the brain region under study^82^.

Predicted EGF signaling was also different in both 5xFAD mice and postmortem human AD samples relative to controls. Samples from control mice and non-cognitively impaired postmortem brain samples exhibited predicted EGF signaling from OPCs to astrocytes (Figure 3G-H). However, no EGF signaling was predicted in 5xFAD mice at 8 months of age, or tissue samples from MCI or AD patients. Notably, there was still predicted EGF signaling in 2-month-old 5xFAD mice, consistent with these animals representing an early disease stage (Figure S6B, E). Regarding the potential function of EGF, a previous study showed that EGF treatment prevented cognitive impairment in female mice, which is consistent with the hypothesis that a loss of EGF signaling contributes to AD^83^. Therefore, our data suggest that signaling through the EGF pathway is reduced in AD, consistent with the protective effect of restoring EGF signaling in diseased brains.

Finally, BMP signaling was also predicted to exhibit similar changes in 5xFAD mice and human AD brains (Figure 3I-J, Figure S6F-G). While no significant BMP signaling was predicted in control or 5xFAD mice at 2 months of age, BMP signaling was predicted to occur at 8 months of age, and slightly more so in controls than 5xFAD mice (Figure S6F-G). The difference was mostly manifest as a reduction in astrocyte-astrocyte signaling during disease progression (Figure S6F-G). Similarly, in human postmortem brain samples, BMP signaling was predicted to be stronger in controls compared to samples of patients with either MCI or AD (Figure 3I-J, S6B). In this case, changes in astrocyte-astrocyte signaling were still predicted, though the most substantial change was predicted to occur in OPCs, in which predicted signaling was reduced in AD (Figure 3I-J). Genes implicated in these changes include *Bmpr2*, *Bmpr1b*, *Bmpr1a*, and *Bmp7* (Figure S6H). While BMP signaling genes *Bmp4* and *Bmp6* have previously been linked to AD and were shown to be upregulated in other mouse models of AD, expression of these two genes was not significantly different in controls vs. disease samples in our mouse data^84^ (Supplementary Table 1). In the human data, *Bmp7* expression was downregulated in AD compared to control samples in astrocytes (logFC = -0.217, adjusted p-value = 8.94E-78, Supplementary Table 1). Notably, *Bmp7*, in contrast to *Bmp4* and *Bmp6*, has a potentially protective effect in AD^85^. Thus, the reduction in *Bmp7* signaling parallels the observed reduction of EGF signaling in AD, as both are thought to be forms of neurotropic signaling and protective against disease. These results thus point to a potential loss of trophic signaling as a common feature of AD pathogenesis in mice and humans.

In addition to pathways where similar inferences were made between mice and humans, we also detected some pathways that had contrasting predicted changes between species. For example, PTPRM had higher predicted signaling in brains from AD patients but lower in 5xFAD mice, PSAP was higher in AD patients but lower in 5xFAD mice, and PDGF was lower in AD patients and higher in 8-month-old 5xFAD mice relative to controls (Figure S6B). PTPRM has been suggested to play a role synapse formation^86^, and though it has not been widely studied in AD, a previous Genome-Wide Association Study identified two SNPs within the PTPRM gene associated with dementia^87^. As these SNPs would not be present in the 5xFAD model, this may be an explanation for the discrepancy between species. In contrast, PSAP, a lysosomal trafficking protein^88^, has been proposed as a potential biomarker for preclinical AD^89^, and a drug targeting PSAP reduced neuronal loss and decreased inflammation in a mouse disease model^90^. The PDGF signaling pathway has also long been linked to AD^91^, with decreases in PDGF signaling being associated with disruption of the blood brain barrier and eventual neurodegeneration^92^. It is possible that these differences across species reflect differences in disease staging, or they may reflect larger differences in disease between the rodent 5xFAD model and human AD.

### NeuronChat predicts a conserved increase in glutamatergic transmission in the 5xFAD rodent model and human AD

Our CellChat analysis provided a set of inter-cellular signaling processes that change during disease, data which implicated changes in signaling between both neurons and glia. This provides a global picture of how inter-cellular signaling may be disrupted in the ENT during the development of AD. One limitation of the CellChat package and database is that it only considers classic ligand-receptor interactions between cells where both ligand and receptor are expressed proteins. However, most neuron-neuron and some neuron-glia communication occurs via small molecule neurotransmitters such as glutamate and GABA. Therefore, this analysis may miss a major component of disease-related changes that develop during AD. To explore potential changes in neurotransmitter-based communication, we applied the newly developed computational package NeuronChat, which utilizes a similar analysis method to CellChat but is adapted for neurotransmitter-based communication^93^. A focus on disease in neurons is warranted, as of the five hdWGCNA gene modules found to be both highly preserved and of high quality, three were found in neuronal populations, and one other in OPCs which have neuron-like features. This raises the possibility that exploring changes in inter-neuronal signaling in 5xFAD mice may have the most relevance for the human AD brain.

We used NeuronChat to compute all the predicted changes in neurotransmitter-based communication in cells from mice and human postmortem brain samples (Figure 4A-E). Changes were observed in 5xFAD mice: at 2 months of age, there was an increase in signaling to oligodendrocytes and endothelial cells (Figure S6I-J). In 8-month-old 5xFAD mice there was an increase in neurotransmitter-based communications, occurring approximately equally in both GABAergic and glutamatergic cells (Figure 4A-B, E). In human postmortem AD brain samples, most changes in signaling were predicted to occur from glutamatergic neurons, with a general increase in predicted communication from glutamatergic neurons to both GABA and glutamatergic neurons in the AD brain (Figure 4C-D). Several predicted increased interactions were observed in both 5xFAD mice at 8 months of age and human AD, including between glutamate and AMPA receptors (Glu-Gria1, Glu-Gria2, Glu-Gria4), NMDA receptors (Glu-Grin2b and Glu-Grin1), kainate receptors (Glu-Grik2), metabotropic glutamate receptors (Glu-Grm1, Glu-Grm5, Glu-Grm8), and the neuronal adhesion molecules neurexin and neuroligin (Nrxn3_Nlgn1; Figure 4E). Notably, no changes in GABAergic signaling pathways were predicted. In addition, genes mediating interactions – *Gria2, Grin1*, *Grin2b, Grm1*, and *Grm5*w – were also present in the preserved hdWGCNA module Glut-M1 (Supplementary Table 2), indicating that expression of genes for these receptors varied across cells in a coordinated fashion across species. When considering genes implicated in glutamate production rather than receptors, we found that *Gls* was significantly upregulated in glutamatergic neurons in 8-month-old 5xFAD mice and tissue samples from human AD brains (logFC = 0.0594, adjusted p-value = 4.79E-05, and logFC = 0.427, adjusted p-value = 4.49E-14, Supplementary Table 1). *Gls* was downregulated in 2-month-old 5xFAD mice (logFC = -0.177, adjusted p-value = 7.45E-71, Supplementary Table 1), and not significantly altered in MCI. *Slc17a7*, or vesicular glutamate transporter 1 (vGluT1), was also upregulated in glutamatergic neurons from both 8-month-old 5xFAD mice and human AD brain samples (logFC = 0.0999, adjusted p-value = 2.95E-10, and logFC = 0.271, adjusted p-value = 0.0464, Supplementary Table 1). Similarly to *Gls*, it was downregulated in 2-month-old 5xFAD mice (logFC = - 0.0777, adjusted p-value = 5.46E-13, Supplementary Table 1), and not significantly differentially expressed in MCI.

**Figure 4:**
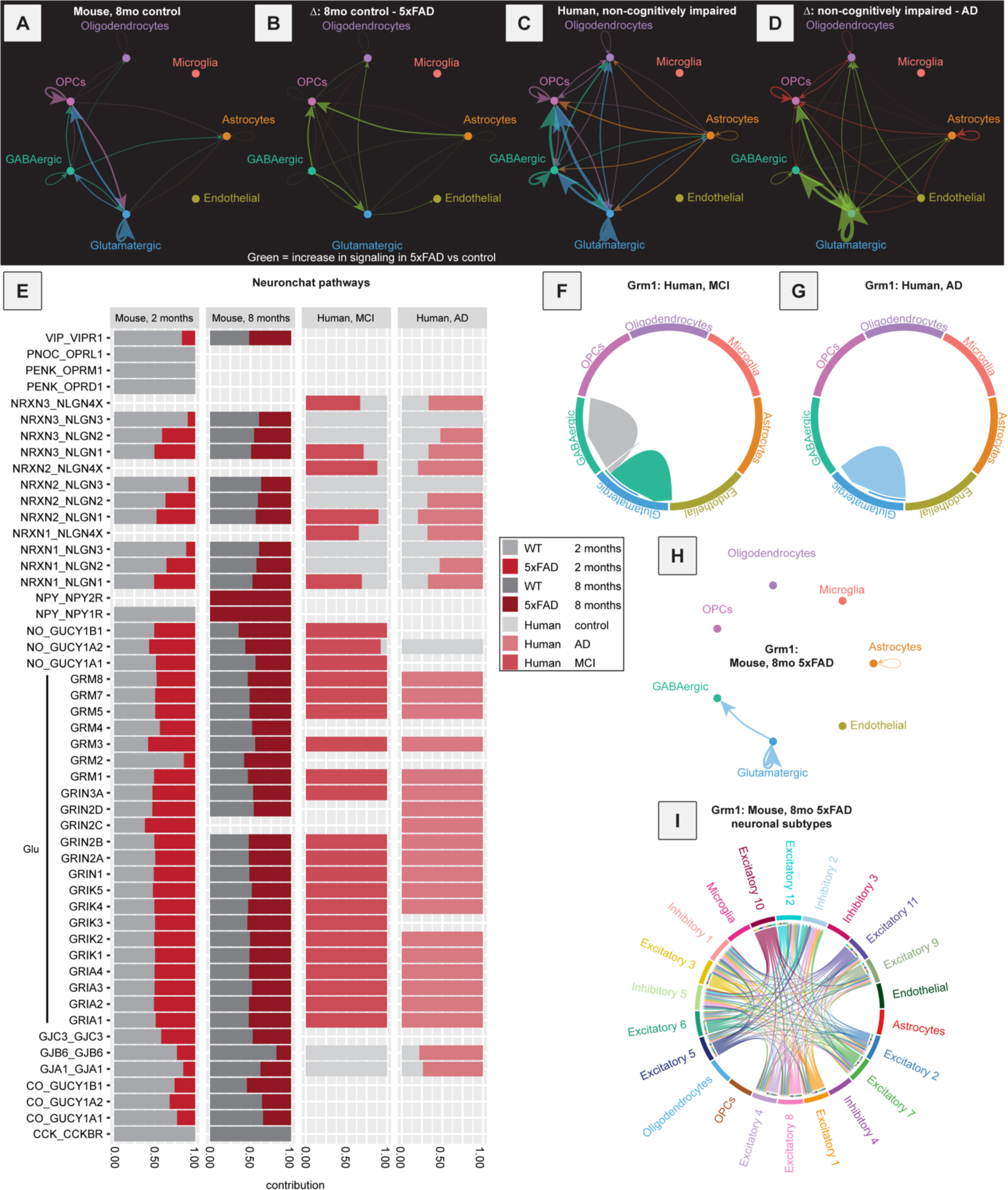
NeuronChat analysis of mouse and human data reveal an elevation of glutamatergic signaling during AD progression. **A**, Signaling inferred for all NeuronChat pathways detected in 8-month-old control mice, aggregated together by signaling strength. **B**, Differences in signaling inferred for all NeuronChat pathways aggregated together by signaling strength in 5xFAD versus control mice at 8 months of age. An increase in signaling in 5xFAD vs control is indicated in green, and a decrease in signaling in red. **C**, Signaling inferred for all NeuronChat pathways detected in samples from non-cognitively impaired human samples, aggregated together by signaling strength. **D**, Differences in signaling inferred for all NeuronChat pathways aggregated together by signaling strength in AD versus non-cognitively impaired individuals. An increase in signaling in AD vs control is indicated in green, and a decrease in signaling in red. **E**, Comparison of predicted signaling for all NeuronChat pathways in 5xFAD versus control mice at 2 and 8 months of age (left), and in samples from individuals with MCI or AD versus those without cognitive impairment (right) **F**, Signaling inferred for *Grm1* in MCI between different cell types. **G**, Signaling inferred via *Grm1* in AD. **H**, Signaling inferred for *Grm1* in 8-month-old 5xFAD mice. **I**, Same data as (**H**), broken down by individual neuronal subtypes.

Related to increased glutamatergic signaling in AD, there is a long-held hypothesis in AD that excitotoxicity mediated by excessive glutamatergic signaling is a key contributor to disease pathogenesis^94–96^. Our combined observations that 1) most predicted changes in neurotransmitter-based interactions occurred via glutamatergic signaling, 2) glutamatergic neurons had two gene expression modules that were both high quality and highly preserved, more than any other cell type, 3) the EPHA signaling pathway is predicted to strongly impact glutamatergic neurons all support this hypothesis. As many of the genes relevant to glutamatergic signaling (*Gria2, Grin1*, *Grin2b, Grm1*, and *Grm5*) were part of Glut-M1 as identified by hdWGCNA, we were curious what other genes may also be in this module. As expected from the GO analysis shown in Figure 2J-L, many ion channel genes were found that contribute to intrinsic neuronal excitability, including sodium ion channel genes *Scn1a*, *Scn2a*, *Scn3a*, *Scn3b*, *Scn8a*, and calcium ion channel genes *Cacna1e*, *Cacna1b*, *Cacna1c*, *Cacna2d1*, and *Cacng3* (Supplementary Table 2). Of these genes, *Cacna1b* and *Cacna1c* were significantly upregulated in human AD, and *Cacna1c* was significantly upregulated in 5xFAD at 8 months of age (Supplementary Table 1), positioning this gene to play a conserved and potentially key role in AD. Upregulation of these genes would likely lead to an elevation in intrinsic neuronal excitability, though it is not clear if these changes may be drivers of pathogenesis, or compensatory responses to disease development. *Cacna1c* has previously been linked to bipolar disorder and depression^97^, and more recently to AD using exploratory bioinformatic analyses^98^. In mice, an increase in *Cacna1c* expression has been negatively correlated with object recognition memory^99^. Interestingly, its expression in reactive astrocytes has also been associated with plaque formation, providing a potential mechanistic role for *Cacna1c* in AD pathogenesis^100^.

We next looked more closely at one particular pathway, the glutamate to metabotropic glutamate receptor 1 (Glu-Grm1, a Gq-coupled GPCR that activates phospholipase C), because 1) Glu-Grm1 was within the highly conserved and high-quality Glut-M1 module, 2) it was identified as differentially regulated in NeuronChat analyses, and 3) it was upregulated in glutamatergic neurons in 5xFAD mice at 8 months of age (logFC = 0.161, adjusted p-value = 4.49E-14, Supplementary Table 1). Unsurprisingly, in both 8-month-old 5xFAD mice and human postmortem brain samples from patients with MCI or AD, glutamatergic neurons were the predicted sender of this Glu-Grm1 interaction in all cases (Figure 4F-H). To define which neuron types may be involved in the predicted Glu-Grm1 interaction, we examined communication in specific annotated neuronal cell clusters of 8-month-old 5xFAD mice (Figure 4I). Every excitatory cluster (subtypes 1-12) exhibited at least some predicted outbound communication via Glu-Grm1, as expected, and inbound glutamate was also widespread, except for in subtype 10 in both 5xFAD and control mice. Interestingly, for GABAergic interneurons, inbound Glu-Grm1 interactions were only predicted onto some subtypes; for example, GABAergic subclusters inhibitory 3, characterized by *Lamp5* expression (Figure S2C), and inhibitory 4, characterized by *Pvalb* expression (Figure S2C), did not have any significant predicted Grm1-Glu communication (Figure 4I), which reflects low expression of *Grm1* in these subclusters. These results suggest that elevations in glutamatergic signaling may differentially impact select subtypes of glutamatergic and GABAergic inhibitory neurons in the ENT.

In addition to glutamatergic signaling, we also found differences in predicted interactions via neuronal cell adhesion molecules. For example, we observed in NeuronChat analyses a predicted increase in communication via Neurexin-3 and Neuroligin-1 in human MCI and AD, though this was not seen in mice (Nrxn3-Nlgn1, Figure 4E). Similar to what we found in data from human postmortem brain samples, expression of *Nlgn1* was significantly upregulated in glutamatergic neurons in 5xFAD mice at 8 months of age (log2FC = 0.163, adjusted p-value = 7.66E-16, Supplementary Table 1), and *Nrxn3* was upregulated in astrocytes, though this was not statistically significant (log2FC = 0.148, adjusted p-value = 0.357, Supplementary Table 1). Neuroligins are post-synaptic cell-adhesion molecules on excitatory synapses^101^, and expression of *Nlgn1* has previously been shown to induce increases in glutamatergic synapse density and activity^102^. We also observed an increase in the expression of neuroligin-related binding proteins in the GO term from GluT M1 (Figure 2J). Thus, the observed upregulation of *Nlgn1* is consistent with an elevation in excitatory synaptic communication, which would be expected to contribute to excitotoxicity. *Nrxn3* is a pre-synaptic protein which can bind to neuroligins, among other proteins, and has been found to be downregulated in postmortem brain samples from AD patients, though these samples were from the hippocampus^103^. Indeed, the function of *Nrxn3* is likely brain region-dependent^104^. Taken together, our results indicate that increased glutamatergic signaling in AD occurs via changes in multiple cell types including glutamatergic neurons but also GABAergic neurons, OPCs, and astrocytes, consistent with a system-wide disease state.

### Disease-associated epigenetic regulation reflects an elevation in glutamatergic communication

The observed systems-wide remodeling of communication between both neurons and glia in the brain are likely dependent on epigenetic modifications that occur during disease. In our rodent sequencing experiments, we performed a multiomic assessment (combined snRNA-seq and snATAC-seq) which enables us to link chromatin accessibility to gene expression across all cell types.

To assess the relationship between chromatin accessibility and gene expression, we utilized DIRECT-NET, which identifies both cis-regulatory and trans-regulatory elements within cellular DNA. Cis-regulatory elements are detected using a machine learning method that links peaks within the snATAC-seq data to gene expression values reflected in the snRNA-seq data, whereas trans-regulatory elements are detected by matching sequences within peaks to known transcription factors. DIRECT-NET was run on data from 5xFAD and control mice from 2-month-old and 8-month-old groups, run separately. This was done so that if links between peaks to specific genes were only found in one genotype or age, we could isolate in which conditions these links were identified. We focused our initial analyses on genes identified in our NeuronChat analyses. Results showed some links specific to 5xFAD, some specific to control mice, and some present in both genotypes (Figure 5A-C). We found that genes encoding glutamate receptors such as *Gria4* and *Grin2b* had many high confidence (HC) links to peaks in the ATAC-seq data, but none in coding regions of the genome outside of the gene of interest itself (Figure 5A-B). In contrast, *Gria4* had 2 HC links outside of the *Gria4* gene locus, which may indicate that these regions are important in the expression of *Gria4*, though they were not specific to either WT or 5xFAD mice, so we did not pursue these regions further. *Grin2b* had more links to loci external to coding regions with some being specific to 5xFAD or control mice.

**Figure 5:**
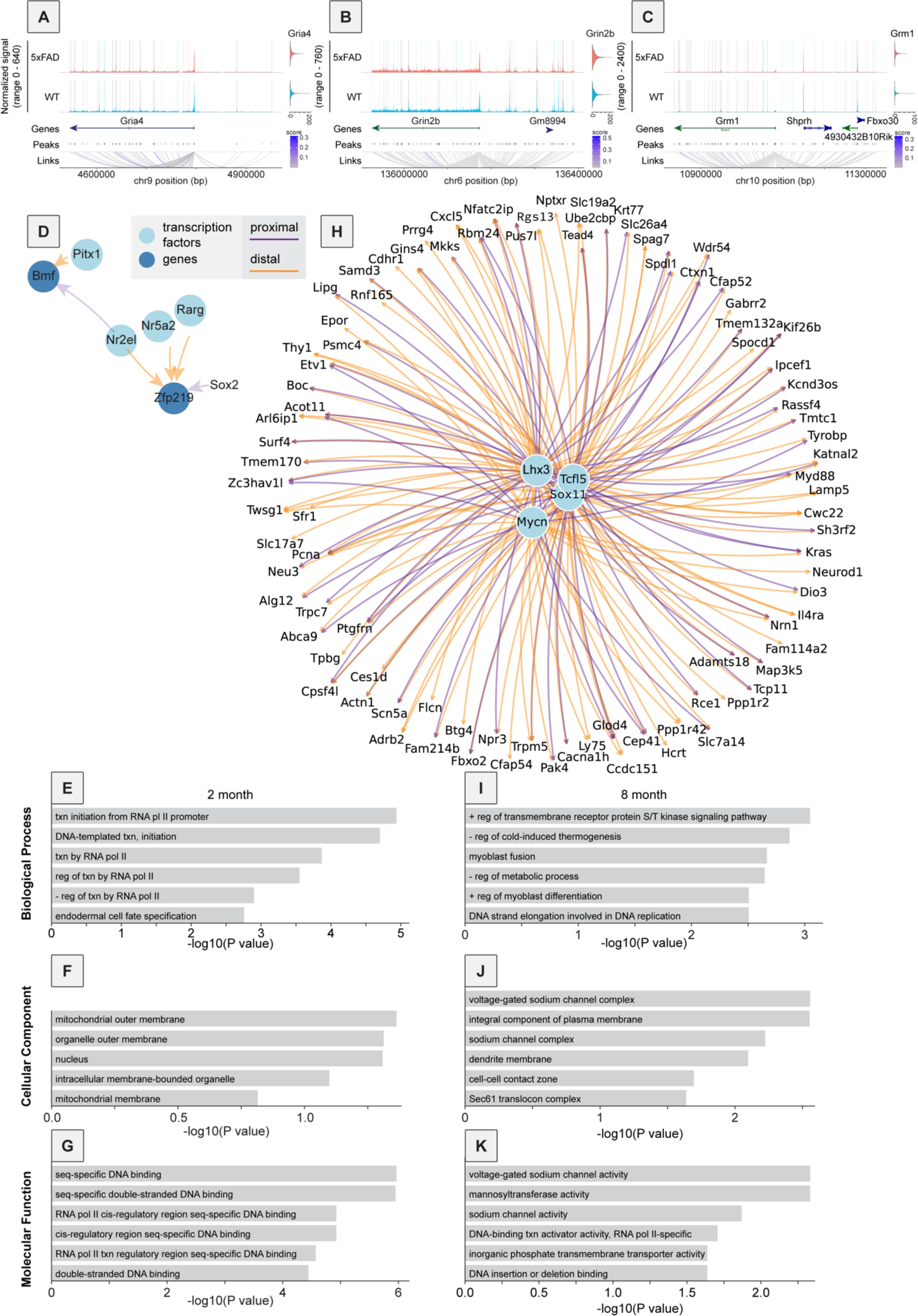
Multiomics approach reveals links between glutamatergic signaling via *Grm1* and trophic signaling via BMPs. **A-C**, Coverage plots of mouse ATAC-seq data with peaks highlighted according to what genotype linkages were detected using DIRECT-NET. Blue indicates linkage to the gene of interest in control mice only. Red indicates linkage to the gene of interest in 5xFAD mice only. Purple indicates linkage to the gene of interest in both genotypes. **A**, Coverage plot of *Gria4* and the surrounding genomic area with links to the *Gria4* transcriptional start site (TSS). A violin plot of *Gria4* RNA expression is shown to the right. **B**, Coverage plot of *Grin2b* and surrounding genomic area with links to the *Grin2b* TSS. **C**, Coverage plot of *Grm1* and surrounding genomic area with links to the *Grm1* TSS. **D**, GRN resulting from DIRECT-NET analysis of genes upregulated in 5xFAD mice at 2 months of age. **E-G**, GO analysis of genes in the GRN obtained in Figure 5D for biological processes (E), cellular component (F), and molecular function (G). **H**, GRN resulting from DIRECT-NET analysis of genes upregulated in 5xFAD mice at 8 months of age. **I-K**, GO analysis of genes in the GRN obtained in Figure 5H for biological processes (I), cellular component (J), and molecular function (K).

Of particular interest was *Grm1*, as in mice at 8 months of age, *Grm1* was upregulated in 5xFAD mice relative to controls (logFC = 0.161, adjusted p-value = 4.49E-14, Supplementary Table 1) and had multiple linked peaks specific to each genotype. In addition to non-coding loci, the expression of this gene was linked to expression of multiple other genes. For example, we observed a link specific in 5xFAD mice from *Grm1* to the transcription start site of the gene *Shprh* (Figure 5C). To our knowledge, *Shprh* has not previously been linked to AD, though it is known to be highly expressed in the human brain and a potential tumor suppressor gene^105^. There was also a link to the transcription start site of the E3 ubiquitin ligase Fbxo30, which was specific to control mice and not observed in 5xFAD mice. Interestingly, Fbxo30 has previously been implicated in the positive regulation of BMP signaling^106^, which we noted in our CellChat analyses in Figure 3 and S6. While Glu-Grm1 interactions identified by NeuronChat were limited to glutamatergic neurons and certain GABAergic neurons, changes in BMP signaling were predicted to occur between astrocytes and other cell types (e.g., Figure 3I-3J, S6F-G). This finding linking *Grm1* expression to genes that control BMP signaling suggests that this may be another route, albeit indirect, by which neurons and glial cells may interact.

We next proceeded to expand our multiomic assessment to any gene that was both upregulated in 5xFAD mice according to our snRNA-seq data, and any loci differentially accessible according to our snATAC-seq data. In 2-month-old mice, DIRECT-NET analyses resulted in a small gene regulatory network (GRN) which included transcription factors Zfp219 and Bmf (Figure 5D). This network’s small size was largely because at 2 months of age, very few loci were significantly differentially accessible between control and 5xFAD mice, which is expected given the early time point in disease progression. To characterize genes within the network we performed a GO analysis and found, unsurprisingly given the role of Zfp219 as a transcription factor, that genes in the network were highly associated with transcription and DNA binding (Figure 5E-G).

Results from this same analysis method but at 8 months of age yielded a much larger GRN (Figure 5H), consistent with the later disease stage, resulting in larger and more substantial changes in chromatin accessibility and gene expression regulation. Overall, this GRN contained 4 transcription factors and 89 genes. GO analysis on this set of genes showed not only association with regulation of transcription and DNA binding, as expected, but also association with ion channel activity (Figure 5I-K). Specifically, voltage-gated sodium channel activity and sodium channel activity were among the top 3 terms relating to molecular function (Figure 5K), another indication that regulation of ion channels may be a principal feature of disease, both at the gene expression and chromatin regulation levels. Notably, one of these transcription factors within the network, Sox11, has previously been implicated in controlling intrinsic neuronal excitability, with its expression being activity-regulated^107^, raising the possibility that this gene may play an important role in regulating the elevation of intrinsic neuronal excitability that occurs in AD.

## DISCUSSION

Here, we show that while inferred signaling changes during AD are widespread between different cell types in the ENT in both 5xFAD mice and human AD, changes in glutamatergic signaling are the most prominent and shared feature of disease in mice and postmortem human samples. This conclusion is supported by several lines of evidence. First, two of the five total gene expression modules identified using hdWGCNA that were both highly conserved and high-quality were found in glutamatergic neurons, while a third was found in GABAergic neurons, and a fourth in OPCs, which all have processes related to neuronal excitability and/or neuronal synapses (Figure 2, S5). In addition, we found using CellChat that the major signaling changes occurred in, glutamatergic cells, GABAergic cells, and OPCs both early and late in disease (Figure 3). Changes in predicted EPHA signaling in AD mostly occurred in glutamatergic neurons (Figure 3D), but multiple genes associated with EPHA signaling were also observed in the OPC gene expression module M1, which also contained the GO terms cAMP- and cGMP-binding, which is critical for EPHA signaling (Figure 2G, Supplementary Table 2). Lastly, our analysis focusing on neuron-specific modes of communication using NeuronChat indicated an elevation in glutamatergic signaling as the main feature of AD pathogenesis in both mice and humans, including the up-regulation of *Slc17a7*, *Gls*, and *Grm1* (Figure 4, Supplementary Table 1). Regulation of *Grm1* was linked to *Fbxo30*, which in turn regulates BMP signaling. As both *Grm1* and BMP signaling processes are reduced in AD, this may be a mechanism by which changes in glutamatergic signaling coordinate with other cell populations to trigger wider modifications in signaling between cell types.

### CellChat and NeuronChat implicate tissue-wide changes in cellular communication during AD

The advent of single cell RNA sequencing technologies has enabled an unprecedented definition of the molecular changes that occur in a variety of disease processes, including in AD. Indeed, there has been a substantial increase in the number of recent publications detailing the molecular changes that occur during AD in a variety of rodent models and human postmortem tissue samples, at multiple time points and in multiple brain regions^5,6,8,26,108–112^. However, one core limitation of these technologies is that they only explore cell-autonomous changes that occur during disease, and do not consider interactions between cells. Thus, traditional sequencing analyses provide only a partial picture of the disease, making it difficult to explore tissue- and systems-level questions. A previous study utilized CellChat to explore the changes in cellular communication that occur between non-neuronal cells^113^; however, how this relates to neuron-neuron and neuron-glia interactions is only just starting to be characterized in humans^114^, and cross-species comparisons are yet to explored. Here we define the gene expression changes in putative receptor-ligand interactions that are conserved between mouse and humans, and focus on neuron-specific changes in communication, identifying changes in glutamatergic signaling across multiple cell types and signaling pathways. For example, the major cell types that are predicted to engage in inter-cellular signaling are glutamatergic neurons, GABAergic neurons, and OPCs (Figure 3A-C). Given that these cells together contain 4 of the 5 gene expression modules that are well-preserved and high-quality between mouse and human, these findings further support the need to understand how changes in communication between these cells contribute to disease progression. In the future, our findings will be further enhanced by the application of spatial transcriptomic technologies, which will allow us to add a spatial dimension to the changes in cellular communication that we observed here. We have recently developed several computational packages designed to integrate snRNA-seq and spatial transcriptomic datasets, including COMMOT^115^, which should further allow us to link changes in gene expression to spatial patterns of cellular communication, as well as pathology *in situ* including amyloid plaques and tau tangles.

One of the common changes between the 5xFAD model and postmortem human AD brains was decreased BMP signaling. This was seen early in the 5xFAD mouse model at 2 months of age and continued through 8 months. In data from human postmortem samples, expression was reduced in both MCI and AD. *Bmp7*, which has not been the focus of many previous studies in the context of AD^116^, stood out as being both highly expressed and significantly differentially expressed across conditions, particularly in astrocytes and OPCs (Figure S6H, Supplementary table 1). BMP signaling has many previously established functions, including neurogenesis and astrogliogenesis^116^, and *Bmp6* has even been shown to localize to amyloid-β plaques in the hippocampus (*Bmp7* was not tested)^117^. *Bmp7* specifically has been suggested to be neuroprotective, but of particular interest to us, it has been shown to reduce the negative effect of excess glutamate^118^. The fact that it is downregulated early in 5xFAD mice and in human AD suggests a reduced ability to respond to increased glutamate load, even before disease is severe.

Further cementing the importance of excitotoxicity were our findings regarding the EGF signaling pathway, which was predicted to decrease in 5xFAD samples at 8 months of age compared to controls, and in MCI and AD samples compared to controls. While EGF signaling has a multitude of functions, it, like *Bmp7*, has also been suggested to be neuroprotective, specifically against excess glutamate^119^. Most prominently, findings from NeuronChat indicated that increased glutamatergic signaling was a shared feature between 5xFAD and postmortem AD human tissue samples. While the glutamate receptor *Grm1* was significantly upregulated in 5xFAD mice, this was not the case in AD (Supplementary table 1), despite the NeuronChat results predicting increased signaling via *Grm1*. These observations suggest that perhaps in AD, elevations in the production of glutamate, rather than its receptors, may be driving increased signaling. In addition, we found that the major changes in epigenetic regulation that occur in the 5xFAD model are related to cellular excitability and regulation of synaptic inputs. Indeed, genes that were identified as being part of the disrupted gene regulatory network in 5xFAD mice at 8 months of age largely related to sodium channels, which are a key contributor to excitotoxicity.

### Neuronal excitability and excitotoxicity as the key conserved features of AD in mice and men

We were surprised to find that no gene expression modules in microglia or astrocytes were both well-preserved and high quality between 5xFAD mice and human AD, but that at least one gene expression module from each of the other cell types analyzed with hdWGCNA were significantly different (Figure S4A-C, S5A, S5F, S5I). This means that the coordinated gene expression changes that occur in microglia and astrocytes during disease development in 5xFAD mice are distinct from those that occur in human AD. If this is the case, it means that processes occurring in 5xFAD mice may not translate well to those occurring in humans. However, before making broad conclusions, several limitations to our study should be considered. First, our study only includes samples from the mouse and human ENT; thus, it is not clear if our findings may extend to other brain regions. Second, we only examined two time points in the mouse, at 2 and 8 months of age, and only females. It is possible that if we examined more time points, different time points, or both males and females, that we may observe better conservation in some of these groups. Third, levels of gene expression detected using snRNA-seq may not directly translate to protein levels, and the relationship between RNA and protein may be cell type-dependent.

Despite these limitations, at face value, our results suggest that the set of features of human AD that are best modeled in the 5xFAD model may be changes in neuronal signaling processes and intrinsic excitability, rather than glial cell processes. It is perhaps not surprising that only certain features of human disease are well-modeled in rodents. For example, AD is a complex disease that includes, among other features, progressive aggregation of amyloid and hyperphosphorylated tau. However, most mouse models of AD pathogenesis only model one core feature of AD, for example amyloidogenesis in the 5xFAD mice. Therefore, molecular and cellular responses related to amyloid plaque aggregation and clearance in human AD may be well-recapitulated in the 5xFAD mouse, whereas molecular and cellular responses related to tau may not be. This, however, also raises the more general problem of how useful rodent models of AD are in studying human AD. No rodent model will perfectly capture all the dimensions of human AD, but perhaps we only need them to recapitulate the essential features of AD for the models to be sufficiently useful for therapeutic development. Indeed, the excitotoxicity hypothesis of AD has existed for many decades, though it has somewhat fallen out of favor, replaced in large part by the amyloid hypothesis, as well as involvement of other processes, such as the immune system^120,121^. It is possible that all of these are correct and interrelated; perhaps amyloid deposition triggers engagement of glia and inflammatory processes, which then effect changes in neuronal signaling that then drive excitotoxicity. If this is true, it is even more critically important to understand how changes in inter-cellular communication between glia and neurons may contribute to disease development and severity. Furthermore, it is important to leverage the strengths of the mouse lines that are used, and acknowledge their limitations, so that we can best focus on pathways and features of disease that are conserved between the mouse and human to maximize the translatability of our findings.

## FIGURE LEGENDS

**Supplemental figure 1:**
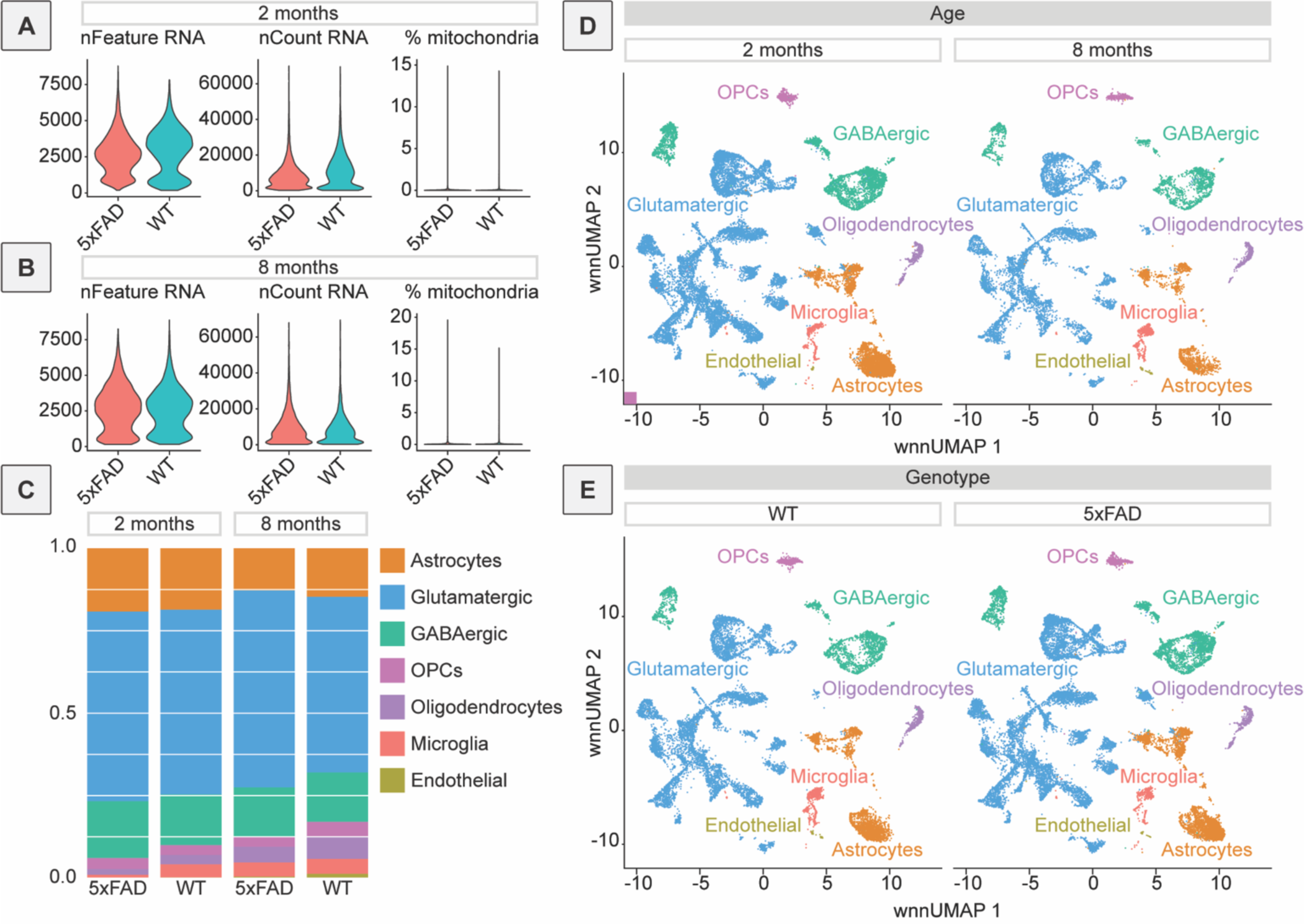
Additional quality control metrics for sequencing data. **A**, Violin plots of QC metrics for snRNA-seq data after filtering out low-quality cells. Data from 5xFAD and control mice at 2 months of age are plotted. Metrics included from left to right are nFeature RNA (number of genes detected), nCount RNA (number of molecules detected), and percent mitochondria (percent mitochondrial genes). **B**, Violin plots of QC metrics for snRNA-seq data after filtering out low-quality cells. Data from 5xFAD and control mice at 8 months of age are plotted. **C**, Bar plots showing cell type proportions for 5xFAD and control mice in the sequencing data. **D**, UMAP visualization of mouse sequencing data, split by age. **E**, UMAP visualization of mouse sequencing data, split by genotype.

**Supplemental Figure 2:**
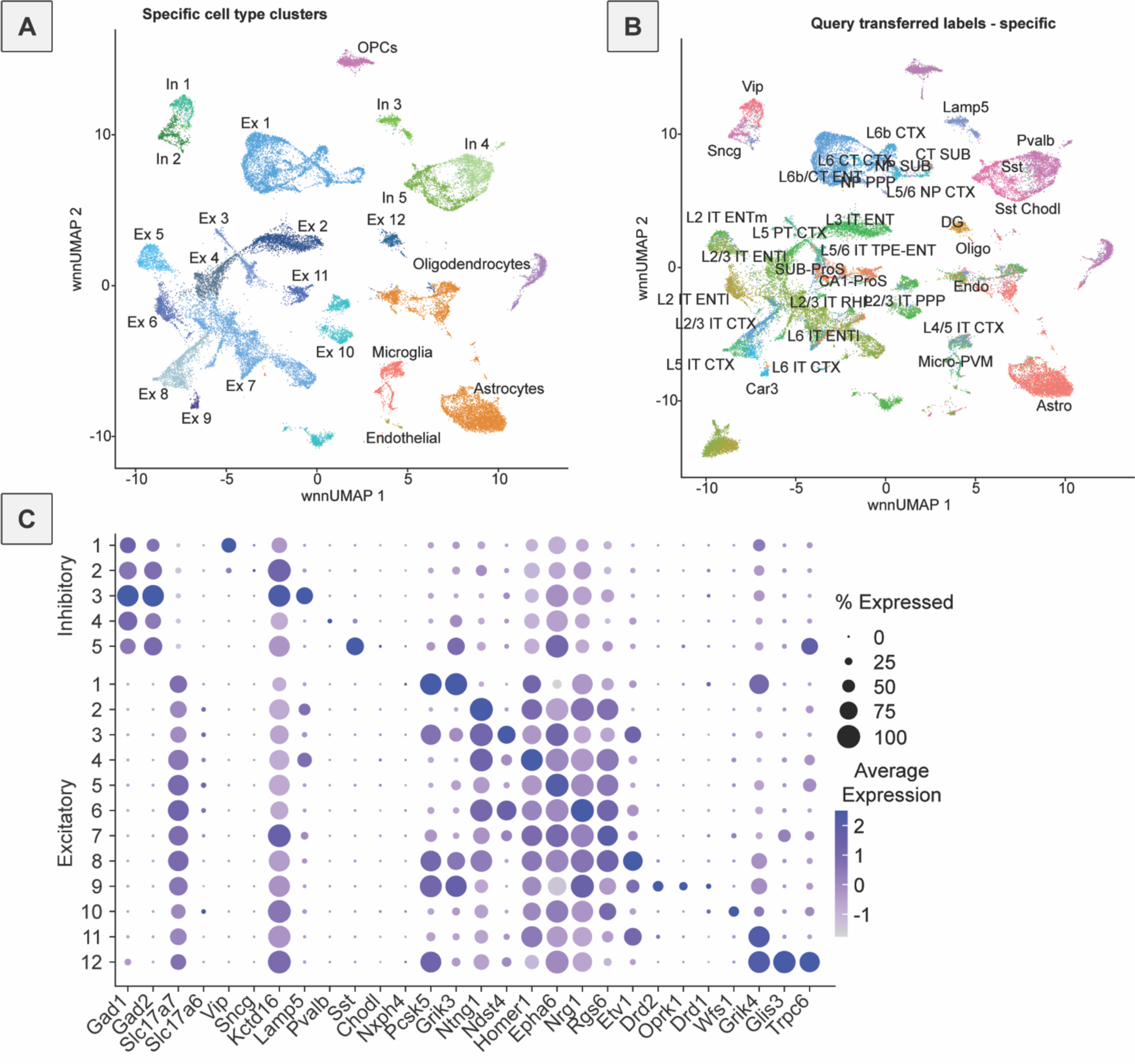
Sub-clustering and further annotation of neurons in 5xFAD and control mouse data. **A**, UMAP visualization of mouse sequencing data with subclusters for different types of GABAergic, or inhibitory (In), neurons and glutamatergic, or excitatory (Ex), neurons. **B**, UMAP visualization of mouse sequencing data annotated based on label transfer from data from the Allen Brain Atlas, including specific neuron types. **C**, Dot plot showing gene expression differences and expression of known marker genes in the subclustered neuronal data as labeled in figure S2A.

**Supplemental Figure 3:**
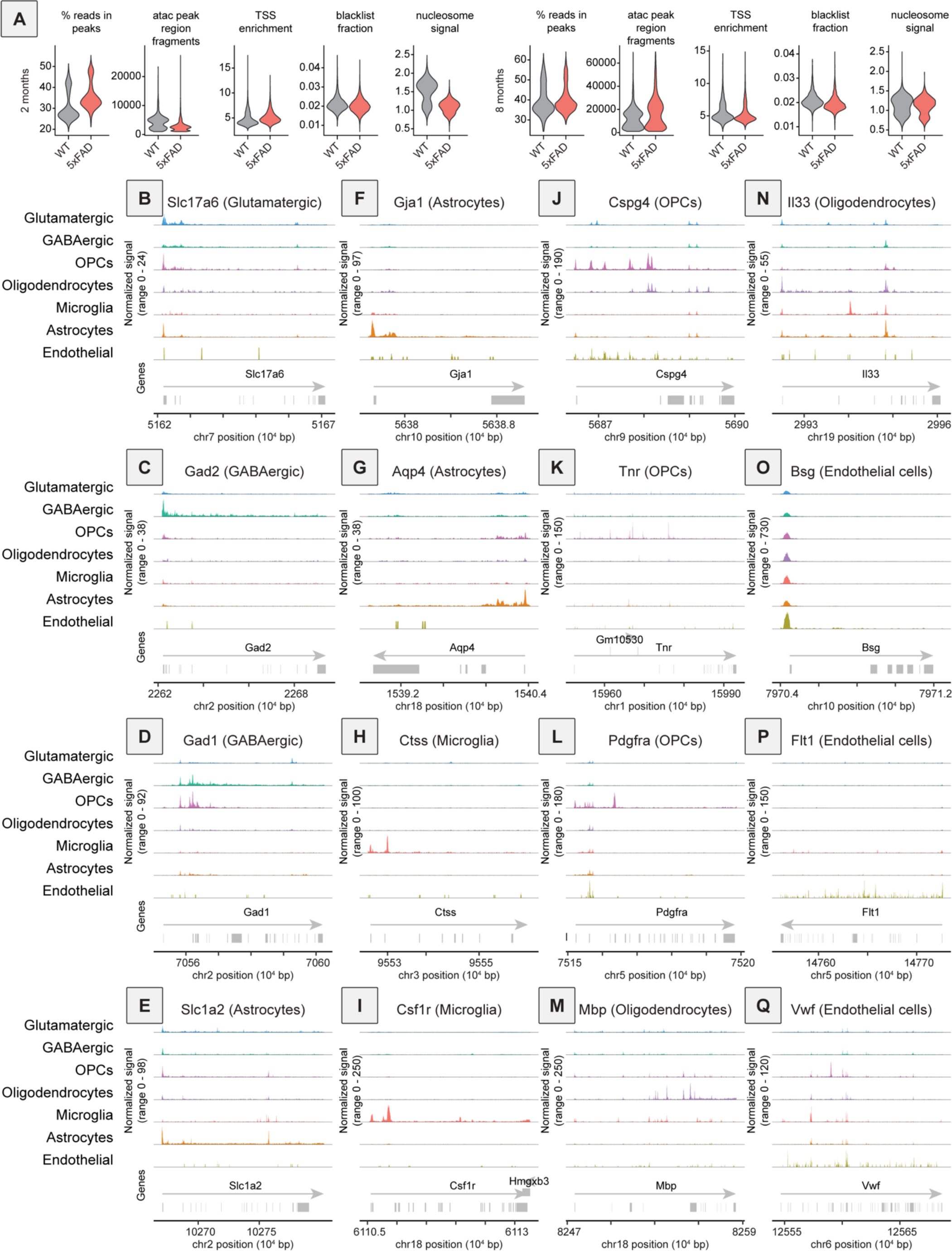
Quality control metrics for snATAC-seq data, and example gene accessibility plots for various cell types in mouse ENT. **A,** Violin plots of QC metrics for snATAC-seq data after filtering out undesirable cells. ATAC QC metrics shown for 5xFAD and control mice at 2 (left) and 8 (right) months of age. From left to right are percent reads in peaks, ATAC peak region fragments, transcription start site enrichment, fraction of reads in blacklist regions, and nucleosome signal. **B,** Coverage plot for *Slc17a6*, a marker gene for glutamatergic neurons. **C,** *Gad2*, a marker gene for GABAergic neurons. **D,** *Gad1*, a marker gene for GABAergic neurons. **E**, *Slc1a2*, a marker gene for astrocytes. **F**, *Gja1*, a marker gene for astrocytes. **G**, *Aqp4*, a marker gene for astrocytes. **H**, *Ctss*, a marker gene for microglia. **I**, *Csf1r*, a marker gene for microglia. **J**, *Cspg4*, a marker gene for OPCs. **K**, *Tnr*, a marker gene for OPCs. **L**, *Pdgfra*, a marker gene for OPCs. **M**, *Mbp*, a marker gene for oligodendrocytes. **N**, *Il33*, a marker gene for oligodendrocytes. **O**, *Bsg*, a marker gene for endothelial cells. **P**, *Flt1*, a marker gene for endothelial cells. **Q**, *Vwf*, a marker gene for endothelial cells.

**Supplemental figure 4:**
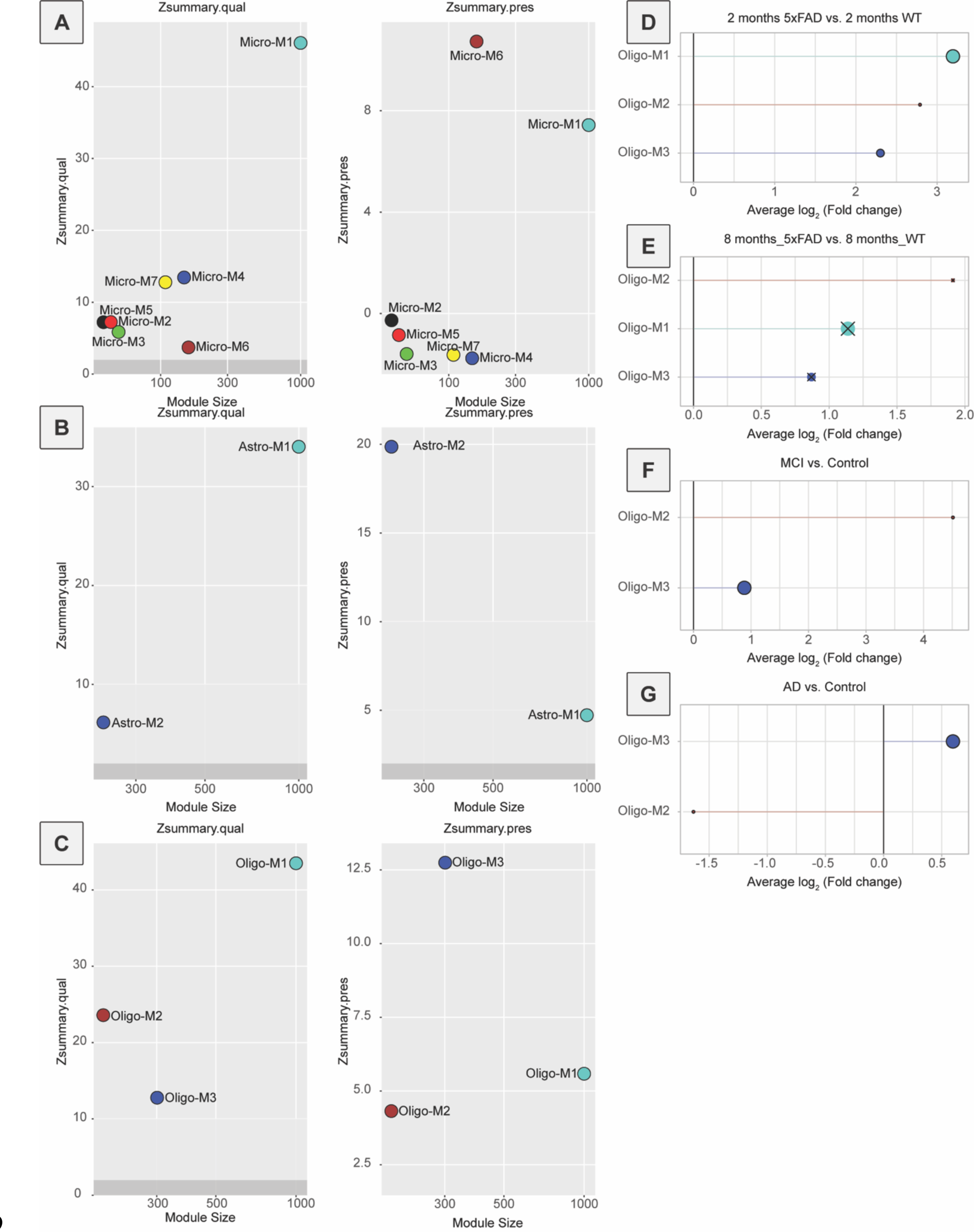
hdWGCNA results for microglia, astrocytes, and oligodendrocytes. **A**, zSummary metrics indicating module quality and preservation across species for modules found within the microglia cluster using hdWGCNA. **B**, zSummary metrics indicating module quality and preservation across species for modules found within the astrocyte cluster using hdWGCNA. **C**, zSummary metrics indicating module quality and preservation across species for modules found within the oligodendrocyte cluster using hdWGCNA. **D**, DME analysis of oligodendrocyte modules in 2-month-old 5xFAD mice. All modules were upregulated in 5xFAD mice at 2 months of age. **E**, DME analysis of oligodendrocyte modules in 8-month-old 5xFAD mice. No modules were significantly different between genotypes at 8 months of age. **F**, DME analysis of oligodendrocyte modules in patients with MCI versus those without cognitive impairment. **G**, DME analysis of oligodendrocyte modules in patients with AD versus those without cognitive impairment.

**Supplemental figure 5:**
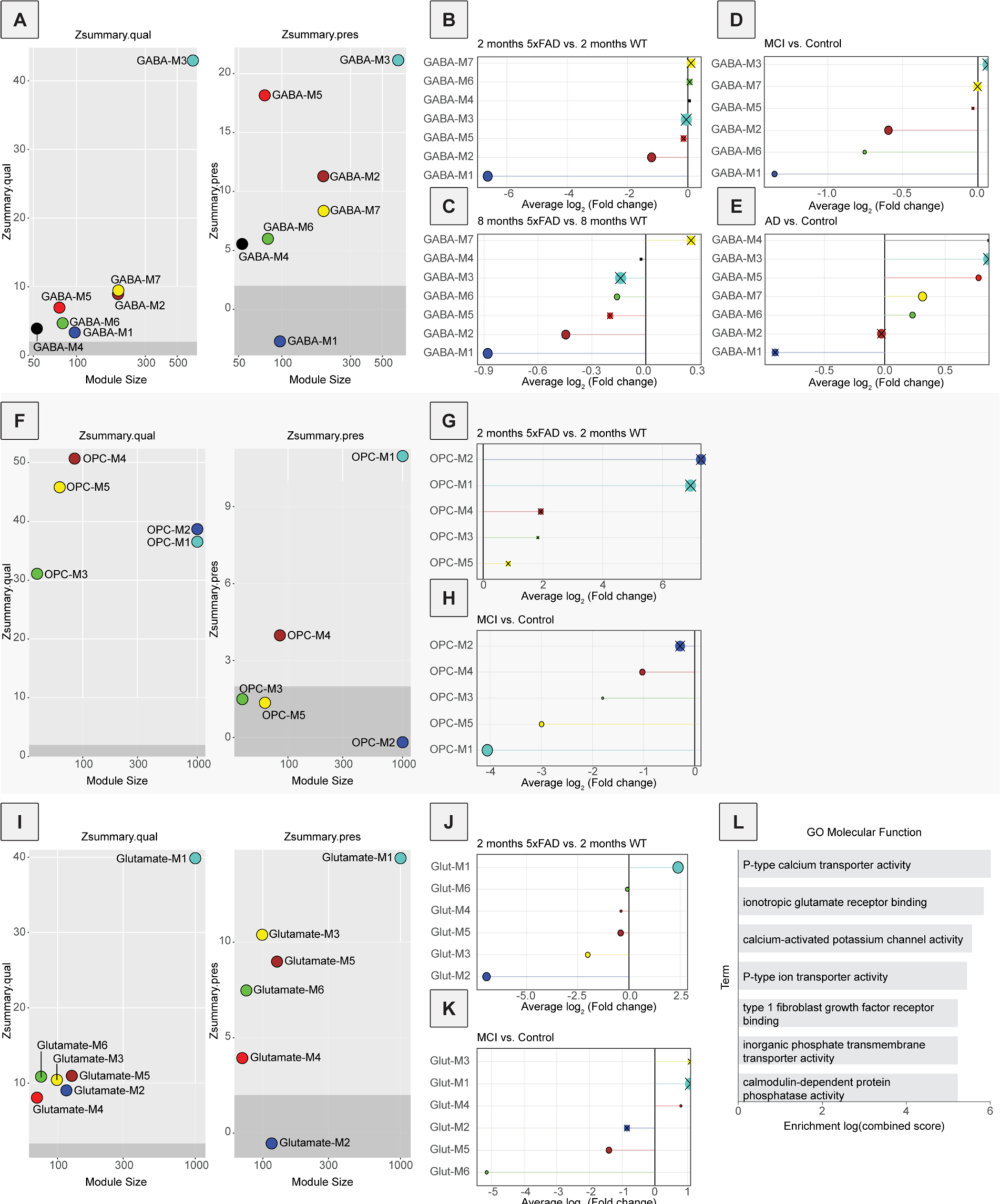
Additional hdWGCNA results for GABAergic neurons, OPCs, and glutamatergic neurons. **A**, zSummary metrics indicating module quality and preservation across species for modules found within the GABAergic neuron cluster using hdWGCNA. **B**, DME analysis of GABAergic neuron modules in 2-month-old 5xFAD mice. **C**, DME analysis of GABAergic neuron modules in 8-month-old 5xFAD mice. **D**, DME analysis of GABAergic neuron modules in MCI. **E**, DME analysis of GABAergic neuron modules in AD. **F**, zSummary metrics indicating module quality and preservation across species for modules found within the OPC cluster using hdWGCNA. **G**, DME analysis of OPC modules in 2-month-old 5xFAD mice. **H**, DME analysis of OPC modules in MCI. **I**, zSummary metrics indicating module quality and preservation across species for modules found within the glutamatergic neuron cluster using hdWGCNA. **J**, DME analysis of glutamatergic neuron modules in 2-month-old 5xFAD mice. **K**, DME analysis of glutamatergic neuron modules in MCI. **L**, GO analysis of genes in the GABA-M3 hdWGCNA module using the molecular function database.

**Supplemental figure 6:**
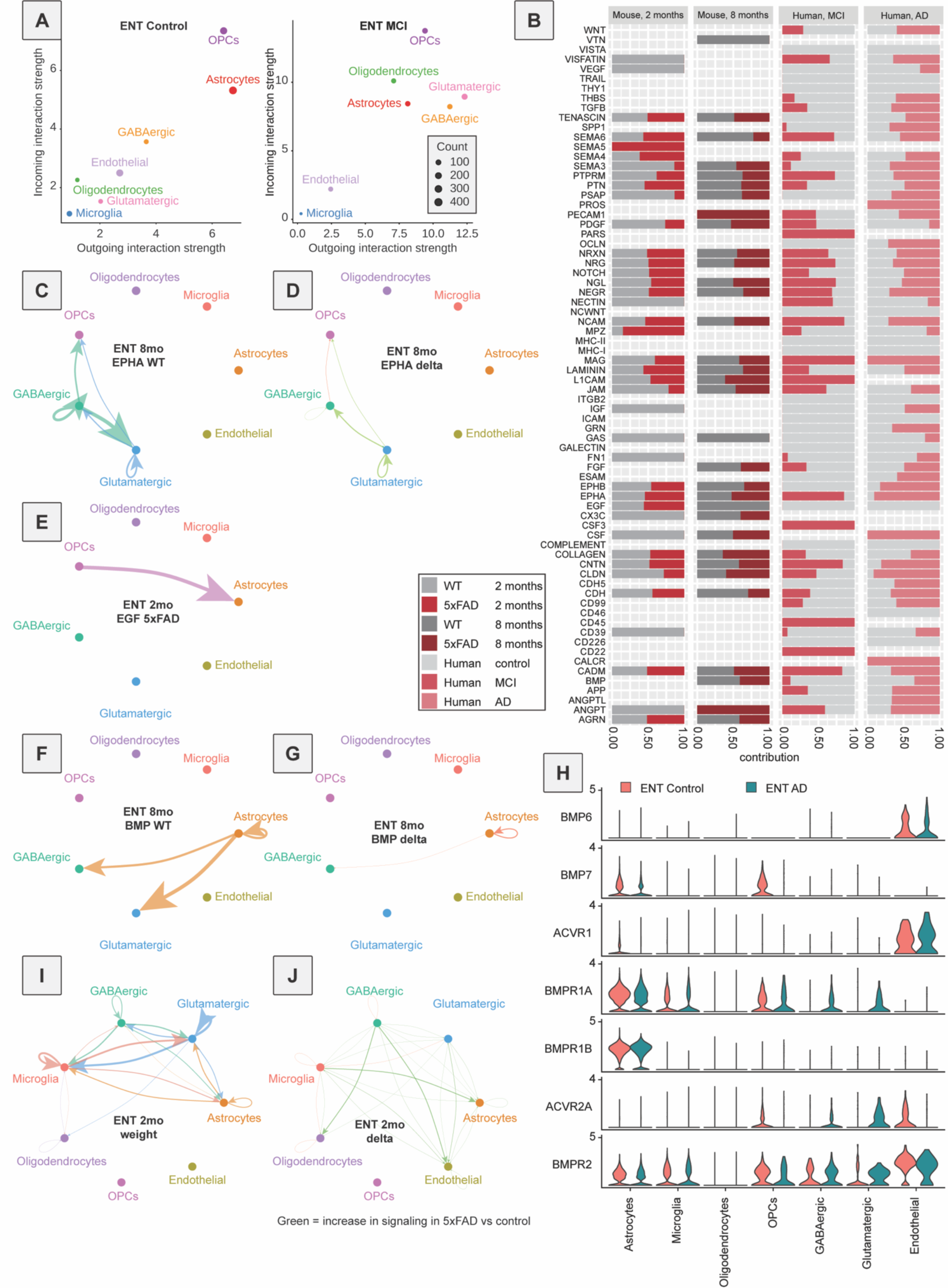
Additional CellChat and NeuronChat results. **A**, Predicted outgoing and incoming cellular signaling with CellChat for each cell type in the ENT of individuals without cognitive impairment (left) and predicted outgoing and incoming cellular signaling with CellChat for each cell type in the ENT of samples from donors with MCI. **B**, Comparison of predicted signaling for all CellChat pathways in 5xFAD versus control mice at 2 and 8 months of age (left), and in individuals with MCI or AD versus those without cognitive impairment (right). **C**, CellChat signaling inferred for EPHA in control mice at 8 months of age. **D**, Differences in CellChat signaling inferred for EPHA in 5xFAD versus control mice at 8 months of age. An increase in signaling in 5xFAD vs control is indicated in green, and a decrease in signaling in red. **E**, CellChat signaling inferred for EGF in 5xFAD mice at 2 months of age. **F**, CellChat signaling inferred for BMP in control mice at 8 months of age. **G**, Differences in CellChat signaling inferred for BMP in 5xFAD versus control mice at 8 months of age. Decreases in signaling are indicated in red. **H**, Violin plots showing expression of genes within the BMP signaling pathway in postmortem human tissue samples from patients with AD and those without cognitive impairment. **I**, NeuronChat signaling inferred for all signaling pathways aggregated in control mice at 2 months of age. **J**, Differences in NeuronChat signaling inferred for all signaling pathways in 5xFAD versus control mice at 2 months of age. An increase in signaling in 5xFAD vs control is indicated in green and a decrease in signaling in red.

**Supplementary Table 1: DGEs.**

**Supplementary Table 2: hdWGCNA modules.**

## Acknowledgements

This work utilized the infrastructure for high-performance and high-throughput computing, research data storage and analysis, and scientific software tool integration built, operated, and updated by the Research Cyberinfrastructure Center (RCIC) at the University of California, Irvine (UCI). The RCIC provides cluster-based systems, application software, and scalable storage to directly support the UCI research community, https://rcic.uci.edu. This work was made possible, in part, through access to the following: the Genomics Research and Technology Hub (formerly Genomics High-Throughput Facility) Shared Resource of the Cancer Center Support Grant (P30CA-062203), the Single Cell Analysis Core shared resource of Complexity, Cooperation and Community in Cancer (U54CA217378), the Genomics-Bioinformatics Core of the Skin Biology Resource Based Center @ UCI (P30AR075047) at the University of California, Irvine and NIH shared instrumentation grants 1S10RR025496-01, 1S10OD010794-01, and 1S10OD021718-01. This work was funded by the Alzheimer’s Association (AARG-NTF-20-685694), New Vision Research (CCAD2020-002), Brightfocus Foundation (A2022031S), NIH (R00DA041445, DP2AG067666, R01NS130044, R01DA056599, 1R01DA054374), Tobacco Related Disease Research Program (T31KT1437, T31P1426), and One Mind (OM-5596678) to KTB, NIH T32GM008620 and NIH F30DA056215 to MH, an NSF grant, DMS1763272, a grant from the Simons Foundation (594598), and NIH T32 GM136624 to KB.

